# Model cyanobacterial consortia reveal a consistent core microbiome independent of inoculation source or cyanobacterial host species

**DOI:** 10.1101/2023.12.09.570939

**Authors:** Andreja Kust, Jackie Zorz, Catalina Cruañas Paniker, Keith Bouma-Gregson, Netravathi Krishnappa, Wendy Liu, Jillian F. Banfield, Spencer Diamond

**Author notes:** **Corresponding author:** Spencer Diamond, Innovative Genomics Institute University of California - Berkeley Innovative Genomics Institute Building, 2151 Berkeley Way, Berkeley, CA 94720. Department of Life Sciences, Imperial College London, London, United Kingdom (Catalina Cruañas Paniker).

## Abstract

Cyanobacteria are integral to biogeochemical cycles, influence climate processes, and hold promise for commercial applications. In natural habitats, they form complex consortia with other microorganisms, where interspecies interactions shape their ecological roles. Although *in vitro* studies of these consortia have significantly advanced our understanding, they often lack the biological replication needed for robust statistical analysis of shared microbiome features and functions. Moreover, the microbiomes of many model cyanobacterial strains, which are central to our understanding of cyanobacterial biology, remain poorly characterized. Here, we expanded on existing *in vitro* approaches by co-culturing five well-characterized model cyanobacterial strains with microorganisms filtered from three distinct freshwater sources, generating 108 stable consortia. Metagenomic analyses revealed that, despite host and inoculum diversity, these consortia converged on a similar set of non-cyanobacterial taxa, forming a 25-species core microbiome. The large number of stable consortia in this study enabled statistical validation of both previously observed and newly identified core microbiome functionalities in micronutrient biosynthesis, metabolite transport, and anoxygenic photosynthesis. Furthermore, core species showed significant enrichment of plasmids, and functions encoded on plasmids suggested plasmid-mediated roles in symbiotic interactions. Overall, our findings uncover the potential microbiomes recruited by key model cyanobacteria, demonstrate that laboratory-enriched consortia retain many taxonomic and functional traits observed more broadly in phototroph-heterotroph assemblages, and show that model cyanobacteria can serve as robust hosts for uncovering functional roles underlying cyanobacterial community dynamics.

## Introduction

In nature cyanobacteria form complex microbial communities [1] where inter-microbial interactions are critical for the resilience, stability, and environmental impact of cyanobacterial assemblages [2–7]. Cyanobacteria-associated microorganisms can act as fixed carbon sinks, modulate cyanobacterial productivity [8], and contribute to the biosynthesis of energetically costly compounds that benefit the entire microbial community [4]. These associated microbes may also play key roles in nutrient remineralization, enhancing the bioavailability of essential nutrients [6], and help mitigate oxidative stress [9]. Conversely, cyanobacteria-associated microorganisms may engage in competitive interactions, including competition with cyanobacteria for inorganic nutrients [9–12]. Cyanobacterial consortia share taxonomic and functional similarities with other phototroph-associated consortia [13] including those of the plant rhizosphere [14, 15], diatoms [16–18] and algae [19], suggesting that common ecological principles drive the assembly of these systems. However, a comprehensive understanding of the dynamics that drive cyanobacterial community assembly, and the roles of core microbial members within these communities, is still evolving [20–26].

Previous studies have provided important insights into cyanobacterial communities using cultivation-independent methods [6, 27–31], synthetic community assemblages [8, 32–35] and isolation and co-cultivation of cyanobacteria with their native microbial partners [21–23, 25, 26, 36–38]. However, *in vitro* consortia, which have made important contributions to our understanding of cyanobacterial communities [8, 21–25, 32–36], often contain few bacterial species and when assembled from isolates can lack taxonomic diversity that accurately reflects natural consortia. Alternatively, naturally sourced communities are not typically cultivated under standardized inoculation conditions or monitored longitudinally. Furthermore, the potential composition and function of microbiomes associated with cyanobacterial model organisms, model species critical to our understanding of cyanobacterial biology, are largely uncharacterized. To address these limitations, we built on previous methodologies to establish a large number of *in vitro* communities grown under standardized laboratory conditions where model cyanobacterial species were used as hosts to enrich co-associated microbiomes from diverse natural inoculation sources. We chose well-characterized model cyanobacterial strains as community hosts including *Synechocystis* spp. [39, 40], *Synechococcus elongatus* [41, 42], and *Nostoc* spp. [43], where information about co-associated communities in natural environments is limited [32, 33, 44]. This system builds upon previous approaches by balancing the simplicity and genetic tractability of model cyanobacteria with the complexity of inoculating these hosts with species derived from environmental samples.

Ultimately this work resulted in the establishment of 108 stable *in vitro* consortia. Using 16S rRNA gene amplicon and genome-resolved metagenomic profiling, we observed that despite the taxonomic composition of the freshwater source microbiomes and diversity of cyanobacterial hosts, the resulting communities rapidly stabilized and became enriched with taxa commonly found in natural freshwater cyanobacterial and other phototrophic symbiotic assemblages [6, 25, 45, 46]. We monitored the stability of these communities over time, including post-cryopreservation revival, confirming their robustness, reproducibility, and ability to be transferred or re-utilized for additional study. The large degree of replication in this study enabled the identification of a 25 species core microbiome, which has not been previously evaluated using *in vitro* efforts. Comparison of core microbiome metagenome assembled genomes (MAGs) to other MAGs recovered in this study confirmed statistically significant enrichment of metabolic traits previously noted observationally in other *in vitro* studies [12, 16, 25, 28, 47] and identified additional functions in porphyrin metabolism and cyanobacterial alkane degradation associated with the core microbiome. Core microbiome members were also significantly enriched in putative plasmids, which encoded functionality potentially supportive of phototroph-heterotroph interactions [48]. Overall, our results uncover the diversity of microbiomes recruited by model cyanobacterial species, demonstrate that laboratory-enriched consortia using model strains retain many of the taxonomic and functional features seen in broader phototroph–heterotroph systems, and establish model cyanobacteria as effective platforms for uncovering new functional roles driving cyanobacterial community dynamics.

## Materials and Methods

### Cyanobacterial selection, inoculation, growth, and passaging

We selected five model cyanobacterial strains—*Synechococcus elongatus* PCC 7942 (7942), *Synechococcus elongatus* UTEX 3055 (3055), *Nostoc* spp., PCC 7120 (A7120), *Synechocystis* spp., PCC 6803 (6803), and *Leptolyngbya* spp., BL0902 (L0902)—based on their genetic tractability, metabolic diversity, and relevance to cyanobacterial biology and biotechnology.

Axenic model cyanobacterial strains were inoculated with bacterial biomass collected from three freshwater bodies (**Table S1**), and were grown in BG-11 [49] medium at 24°C under a 12:12 light cycle and passaged at both weekly and bi-weekly intervals for 12 weeks. Samples were collected at each passage and rigorous standardized assessments of culture contamination, and host dominance were applied to determine if a co-culture should be retained in the study (**Table S2**). For comprehensive details, see **Supplementary Methods** and **Fig. 1A**.

**Figure 1.**
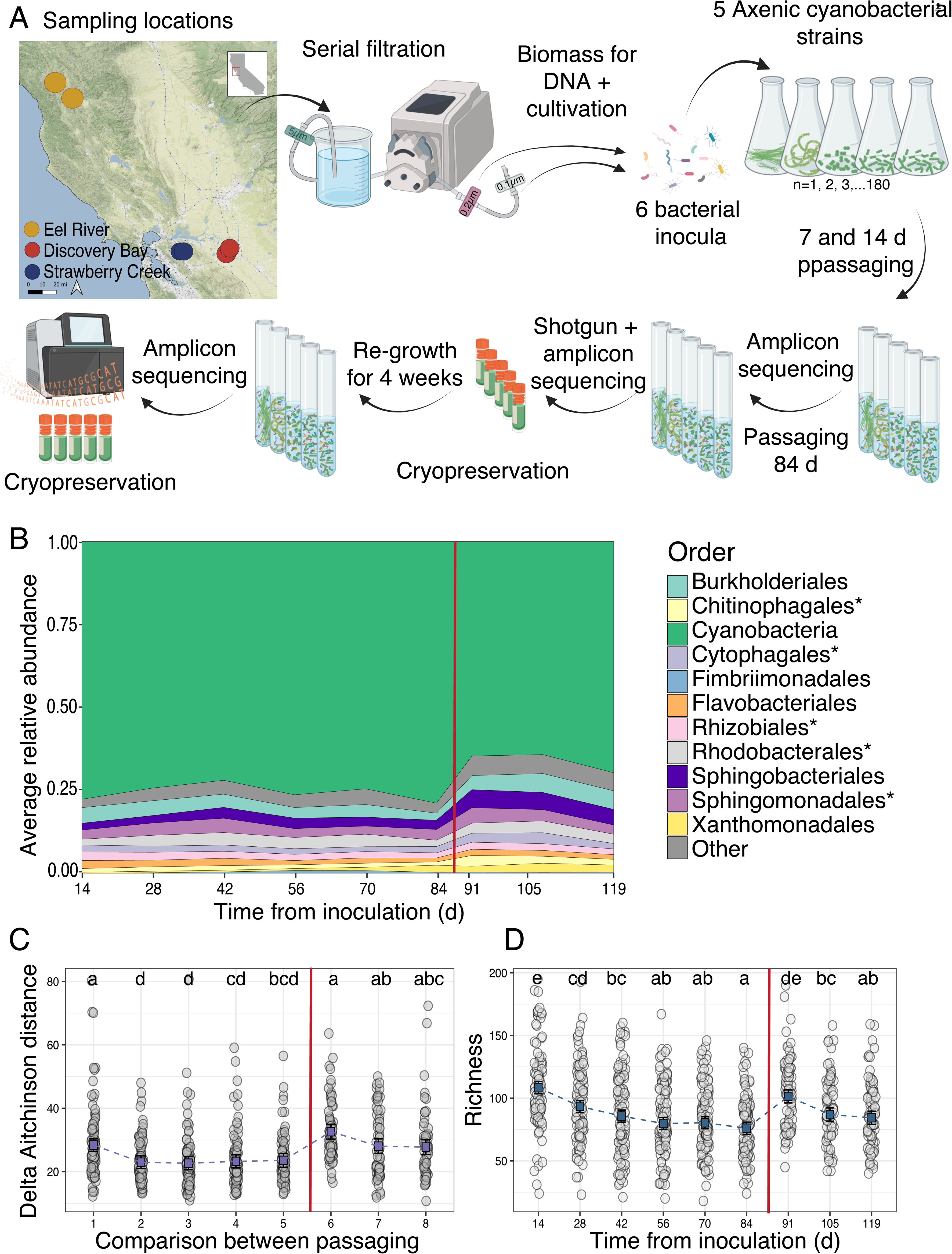
Assessment of *in vitro* community stability over the 119 d passaging experiment. **(A)** Experimental design for cyanobacterial *in vitro* community cultivation. Water samples were collected from three freshwater locations (two sites per location) resulting in six source inocula. Bacterial fractions were prefiltered, then concentrated using 0.2 µm and 0.1 µm filters, combined, and used to inoculate five cyanobacterial host strains in triplicate. Each host was paired with six source inocula twice for passaging at both 7- and 14-day intervals over 12 weeks, resulting in 180 initial cultures. At each passage, samples were collected for microscopic analysis, OD measurements, and 16S rRNA gene amplicon sequencing. After the 84 d passage, samples were collected for metagenomic sequencing and cryopreservation. The frozen consortia were then regrown, passaged, and harvested as before. **(B)** Order-level community composition for *in vitro* communities at 14 d intervals over 119 d, based on 16S rRNA gene amplicon sequencing averaged across all samples. Orders found in the core microbiome are indicated with an asterisk (*). In all plots the vertical red line separates pre-cryopreservation passaging (left) and post-cryopreservation passaging (right). **(C)** Change in Aitchinson distance between previous passage time points for *in vitro* communities at 14 d intervals over 119 d. Gray dots give change for each community compared to itself in the preceding time point, purple squares indicate estimated marginal means, and error bars represent 95% CI of marginal mean. Statistical significance between comparisons estimated using linear mixed effects models with comparisons not sharing the same letter being significantly different (FDR ≤ 0.05). **(D)** Observed richness of *in vitro* communities at 14 d intervals over 119 d (total ASV counts). Gray dots give values for each community at a given time point, blue squares indicate estimated marginal means, and error bars represent 95% CI of marginal mean. Statistical significance between time points estimated using linear mixed effects models with time points not sharing the same letter being significantly different (FDR ≤ 0.05).

### DNA extraction and sequencing

DNA for shotgun metagenomic sequencing was extracted from inoculation source material and biomass of co-culture samples at day 84 of passaging using a modified Qiagen DNeasy PowerSoil Pro Kit protocol. DNA from co-culture samples was sequenced with paired-end reads of 150 bp, whereas DNA from the source material was sequenced with paired-end reads of 250 bp on a NovaSeq System (Illumina). We used longer sequencing reads (250 bp) and a larger library insert size for complex environmental source samples to improve assembly and contig recovery [50].

For 16S rRNA gene amplicon sequencing, genomic DNA was extracted [51] from co-culture samples of all passages. The extracted DNA was amplified for the V4–V5 regions of the 16S rRNA gene using Illumina-compatible primers and sequenced on a MiSeq System (Illumina) with a read depth of 50,000 reads per sample (mean = 46,098 reads per sample). This depth enabled frequent monitoring of community changes and complemented shotgun metagenomic sequencing, which was employed for species-level resolution and functional analysis of gene content. For comprehensive details, see **Supplementary Methods.**

### 16S rRNA gene data processing and statistical analysis

Amplicon sequence variants (ASVs) were generated from fastq files using the USEARCH-UNOISE3 pipeline [52], with modifications, to identify a total of 2,126 ASVs. Taxonomy was assigned using a naive Bayes classifier trained on the SILVA v138 SSU [53] reference database. We ultimately analyzed 869 rRNA gene amplicon samples (**Table S3**) that represented samples collected from all longitudinal passages of co-culture samples determined to be un-contaminanted, dominated by the expected host cyanobacterium, and also analyzed with shotgun genome-resolved metagenomics at day 84. Alpha-diversity (Richness, Shannon) was assessed with rarefied ASV counts (**Table S4**) using linear mixed-effects models. Beta-diversity was analyzed with additive log ratio (ALR) transformed ASVs and Aitchison distances (**Table S5**). Differential abundance was tested with Maaslin2 (**Tables S6**). For comprehensive details, see **Supplementary Methods** and **Code Availability.**

### Assembly independent marker gene analysis of shotgun metagenomic data

As recovering MAGs from complex source samples was challenging, we also employed an analysis where marker genes were directly identified from shotgun metagenomic reads to more accurately compare diversity and taxonomic composition between source samples and co-cultures. This was conducted using ribosomal proteins as in prior work [54]. Using SingleM v0.13.2 [55], unique Ribosomal protein L6 (rpL6) were identified and clustered at 95% nucleotide identity to produce rpL6 OTUs that could be compared across samples (**Table S7**). Over/under-enrichment of bacterial orders in cyanobacterial communities relative to source samples was evaluated with a permutation-based method (**Table S8**). For comprehensive details, see **Supplementary Methods** and **Code Availability**.

### Metagenome assembly, binning, bin de-replication

Metagenome assembly and binning were conducted using methods similar to [50]. However, scaffolds > 5000 bp were reassembled with COBRA v50 [56], and validated circular contigs were re-incorporated into the assembly. Genome binning was performed on a sample by sample basis and only used scaffolds > 2,500 bp using MetaBat2, maxbin2, CONCOCT, and vamb [57–60]. Best MAGs across the four methods for each sample were selected using DasTool [61] and evaluated with checkM [61, 62]. Only MAGs with ≥60% completeness and contamination of ≤10% were retained. MAGs were de-replicated at the species level and representative species MAGs were selected using dRep [63], considering MAGs as the same species if ANI was ≥ 95% across ≥ 10% of the MAG length. This resulted in a non-redundant set of 537 MAGs (**Table S9**). Taxonomic classification was conducted using GTDB-tk [64] classify workflow against the GTDB-R214 taxonomy. For comprehensive details, see **Supplementary Methods.**

### Phylogenetic analysis

GToTree v1.6.34 [65] was employed to detect, align, trim, and concatenate 120 universal single copy genes [66] from non-redundant MAGs as well as an archaeal outgroup species *Acidianus hospitalis* W1 (GCF_000213215.1_2). MAGs with < 10% of markers were excluded. Maximum-likelihood phylogenetic reconstruction was conducted with IQ-TREE version 1.6.12 [67]. The best-fit evolutionary models for each marker protein partition were determined using ModelFinder Plus. Tree topology significance was assessed using ultrafast bootstrap analysis with 1,000 replicates and the Shimodaira-Hasegawa approximate likelihood ratio test (SH-aLRT) with 1,000 replicates. The resulting phylogenetic tree was displayed using iTol v6.8.1 [68]. For comprehensive details, see **Supplementary Methods.**

### MAG abundance mapping and diversity analysis

Shotgun metagenomic reads from each sample were mapped to species representative MAGs with Bowtie2 [69] and coverage quantified with CoverM [70]. MAGs were considered present in a sample if they met stringent coverage and breadth criteria (**Table S10**). Metagenomic samples were removed from analysis when the expected host cyanobacterium comprised ≤ 90 % of the cyanobacterial fraction resulting in 121 shotgun metagenomic samples (108 cyanobacterial cultures and 13 source environment samples) for downstream analysis.

Beta-diversity was assessed using Aitchison distance of MAG mapped read counts, and permutational analysis of variance (PERMANOVA) was used to estimate the influence of sample type (culture or source) and location (**Table S11**). For comprehensive details, see **Supplementary Methods** and **Code Availability.**

### Core microbiome criteria

The concept of a core microbiome lacks a universally accepted definition [71, 72], so we developed an operational framework to identify core microbiome species (MAGs) across co-cultures based on two key criteria: (i) a core species must reproducibly co-occur (was considered present in a co-culture by MAG mapping criteria) with at least four cyanobacterial host species across three independent samples for each host (a minimum of 12 samples); and (ii) it must be consistently recruited (was considered present in a co-culture by MAG mapping criteria) from at least two distinct environmental sources across at least three independent samples per environmental inoculum source (a minimum of 6 samples). This operational definition ensures core species are consistently represented across both host diversity and environmental variability, reducing the likelihood of sampling artifacts or host-specific associations. To asses the relatedness of core microbiome MAGs assigned to the same species, we performed digital DNA-DNA hybridization (dDDH) analysis using the Genome-to-Genome Distance Calculator [73, 74](**Tables S12**).

### Population level single nucleotide variant (SNV) analysis

Population-level SNV analysis was performed on 108 cyanobacterial co-culture samples using inStrain v1.6.4 [75], generating 3,088 pairwise comparisons for 133 MAGs, including 555 comparisons involving core microbiome species. No identical strain sharing was detected across 176 comparisons between geographically distinct inocula (popANI ≥ 99.999%; **Table S13**). Beta regression modeling showed significant effects of inoculum source and host strain on MAG microdiversity (adjusted P ≤ 0.05; **Table S14**). For comprehensive details, see **Supplementary Methods, Supplementary Results,** and **Code Availability.**

### Assessment of co-culture community taxonomic overlap with other studies

We compiled a set of 1,843 MAGs that passed completeness and contamination criteria used in our study (920 distinct species) from 10 genome-resolved studies involving diverse cyanobacterial or plant rhizosphere samples (**Tables S15 and S16**). These included studies of non-axenic cyanobacteria cultures [23, 76–78], environmental cyanobacterial consortia [27, 28, 30, 79], and rhizosphere derived bacteria [80, 81]. Identical species (≥ 95% genome ANI) between the 920 species-representative MAGs from other studies and the 319 MAGs observed in co-cultures of this study were identified using dRep [63]. Additionally we evaluated higher level taxonomic overlap between species-representative MAGs present in our co-cultures and those of other studies at the class and genus level using taxonomic classifications produced using GTDB-tk [64] classify workflow against the GTDB-R214 taxonomy.

### Functional annotation and core microbiome functional enrichment analysis

Predicted proteins from all 537 MAGs were annotated using KOfam HMMs via kofamscan [82], and METABOLIC [83]. Alkane degradation genes were identified using hmmsearch [84] with HMMs from CANT-HYD [85]. To assess core microbiome functional enrichment (n = 25 MAGs) compared to non-core MAGs (n = 512 MAGs), Fisher tests were conducted on KEGG orthologs (KOs) with p-values adjusted for multiple testing (FDR ≤ 0.05). Two tests were performed: one using all MAGs (n = 537 MAGs) and another constrained to Pseudomonadota, Bacteroidota, and Spirochaetota MAGs (n = 391 MAGs) to account for taxonomic bias in the core microbiome MAG set. Broader functional categories were assigned, and differential representation was analyzed using Fisher tests with multiple testing correction. For comprehensive details, see **Tables S17-S20**, **Supplementary Methods,** and **Code Availability**.

### Identification, curation, and analysis of putative mobile genetic elements

Contigs predicted to be mobile genetic elements (MGEs) were identified across 537 MAGs using geNomad [86] filtering for contigs ≥12 kb and FDR ≤ 0.05 (**Table S21**). We evaluated all contigs ≥12 kb that were confidently identified as putative plasmids, given the high frequency of circular element fragmentation in short-read metagenomic assemblies. Plasmid assignment to MAGs was based on a guilt-by-association approach, where plasmids co-binned with a MAG were considered associated. Plasmid content was compared between core, auxiliary, and source genomes using Dunn’s test. All plasmid contigs were aligned to the IMG/PR database [87] using BLASTn for identification (**Table S22**). Genes related to conjugation, mobilization, and replication were identified using HMMs from Conjscan [88] and Pfam [89] via hmmsearch [84]. Plasmids were classified as mobilizable or conjugative based on the presence of key genes (e.g., MOBX, T4CP, virB4). General functional annotation was performed identically as for MAGs detailed above.

For MGE-associated functional enrichment, plasmid-contigs were removed from the 537 MAGs, and KEGG orthology (KO) enrichment analysis was repeated. Differential KO enrichment was compared between the full and MGE-removed analyses using Fisher’s test, with FDR correction (FDR ≤ 0.05) (**Tables S18 and S19**). Fold changes in KO enrichment were calculated between the original and MGE-removed analyses. For comprehensive details, see **Supplementary Methods,** and **Code Availability**.

## Results

### Inoculation of model cyanobacterial strains with bacteria from diverse freshwater sources established stable *in vitro* communities

A total of 108 consortia were retained at the end of 12 weeks of passaging being free of contamination and dominated by the original cyanobacterial host **(Fig. S1 and Supplementary Methods)**. The retention of consortia in the experiment was influenced by the cyanobacterial host, inoculum source, and the interaction between host strain and passage rate. Among the tested strains, 6803, L0902, and A7120 exhibited the highest retention rates (**Supplementary Results**).

Profiling of 16S rRNA gene amplicons showed that despite differences in inoculum source and host strain, communities rapidly stabilized within the first 28 days (**Figs. 1B-D and S2**).

Cryopreservation and revival confirmed the long-term stability of the communities, with diversity metrics and taxonomic composition returning to pre-cryopreservation values (**Fig. 1B-D**). The final consortia set evaluated for this study (84 days post inoculation) closely resembled natural cyanobacterial communities, displaying a consistent taxonomic structure (**Fig. 1, S3, and Supplementary Results**). These results confirm that model cyanobacterial species can form stable, reproducible consortia *in vitro*, simulating natural microbiomes [6, 9, 19, 90].

### Genomic resolution enabled comprehensive characterization of *in vitro* communities

Shotgun sequencing of stable consortia (84 days) and source samples yielded 537 non-redundant species-level MAGs, with 59.8% classified as high-quality (completeness of ≥90%, contamination of ≤5%) and 40.2% as medium-quality (completeness of ≥60%, contamination of ≤10% (**Table S9**). These MAGs, spanning 17 phyla, primarily Bacteroidota and Pseudomonadota (**Fig. 2A**), included 84.2% of MAGs without species-level representatives in existing genomic databases, and59 MAGs unclassified beyond the family level (**Table S9**). The recovered MAGs effectively represented the *in vitro* communities, with 86.2% of co-culture reads mapping back to MAGs compared to only 9.3% of source sample reads mapping back to MAGs on average (**Fig. 2A, Table S2, Supplementary Results**).

**Figure 2.**
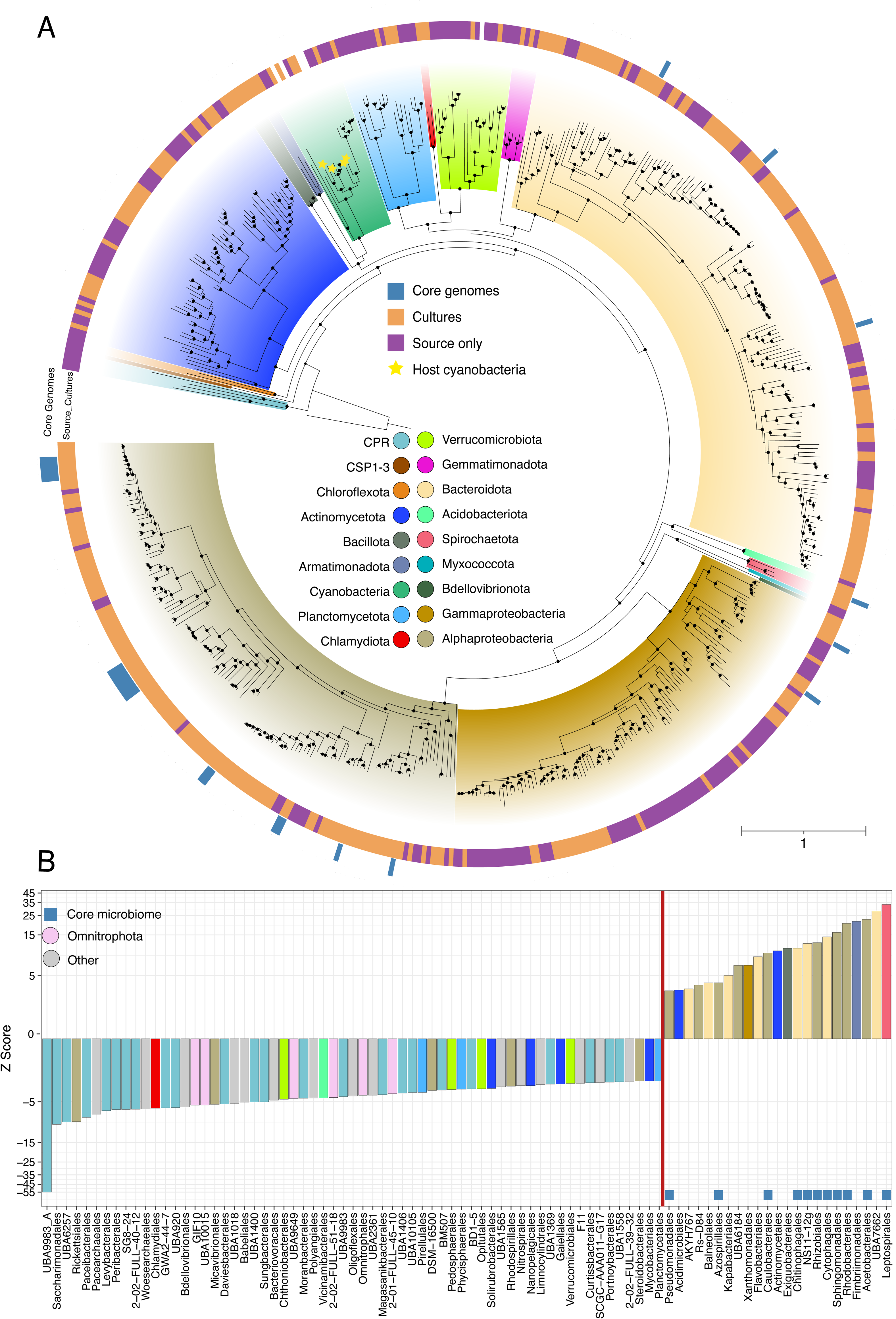
Maximum likelihood phylogenetic tree and enrichment of microbial orders. **(A)** A phylogenetic tree of 537 MAGs in our study constructed using a concatenated alignment of 120 bacterial-specific marker genes. An archaeal reference genome (GCF_000213215.1) served as an outgroup for rooting the tree. Nodes marked with black dots represent a high bootstrap support (≥ 95%, calculated using ufboot with n = 1000 iterations). The outer ring highlights core microbiome MAGs (in blue), and the inner ring differentiates MAGs observed in *in vitro* cultures (orange) and those exclusively observed in source samples (purple). Branch tips with cyanobacterial host species are denoted by yellow stars and have no outer ring colors. Scale bar indicates the number of nucleotide substitutions per site. **(B)** Microbial orders showing significant over or under enrichment when comparing ribosomal protein L6 marker gene OTU (rpL6 OTU) diversity between true *in vitro* community compositions and randomly sampled *in vitro* communities (n = 10,000 permutations) where all observed rpL6 OTUs were sampled. The Z-score indicates the level of enrichment of an order in observed communities relative to communities generated from random sampling. Only orders with statistically significant enrichment values are shown (FDR ≤ 0.05; permutation test). Orders positioned on the left of the red line are underrepresented in true consortia, and those on the right of the line are over-represented in true consortia. Blue squares identify taxa found in the core microbiome. Bars are colored by phylum as indicated in the key of Fig. 2A with additional colors added and noted in the key of Fig. 2B for minor phyla (Other) and Omnitrophota. Also see Tables S8 and S9.

### Specific taxonomic groups become enriched in stable *in vitro* communities

We directly compared diversity between environmental source samples and co-cultures. This analysis was conducted both by comparing MAG diversity (**Fig. S5**) and performing an assembly-independent comparison of ribosomal protein L6 marker OTU diversity (L6OTU; **Fig. 2B and S4**), to overcome low MAG recovery from source samples. Both analyses revealed significantly lower alpha-diversity for *in vitro* consortia relative to source samples (**Fig. S4 and S5**). Using L6OTU analysis we found that 22 orders were overrepresented, whereas 59 were underrepresented within *in vitro* consortia relative to source environments. Bacterial orders from the Pseudomonadota and Bacteroidota were the most frequently enriched in co-cultures, and bacterial orders within the Candidate Phyla Radiation (CPR) and Omnitrophota were underrepresented (**Fig. 2B, Table S8**). Despite differences in inoculum sources, the *in vitro* environment consistently favored the enrichment of specific taxa (**Supplementary Results**).

### Inoculum source and cyanobacterial host strain drive inter-community variation

Stable co-cultures were predominantly composed of their original cyanobacterial host strain, accompanied by prevalent Pseudomonadota and Bacteroidota species, along with less common phyla such as Spirochaetota, Actinomycetota, and Armatimonadota (**Fig. 3A-B and Table S10**). Between co-culture samples, we found that the source of inoculum (*R^2^* = 0.23, *P* < 0.0001; PERMANOVA), the cyanobacterial host strain (*R^2^* = 0.11, *P* < 0.0001; PERMANOVA), and the interaction between the inoculum source and host strain (*R*^2^ = 0.08, *P* < 0.0001; PERMANOVA), all contributed to statistically significant variations in community composition, with inoculum source having the strongest influence (**Fig. S5G-I**). Despite these variations, communities converged toward a broadly similar taxonomic structure with enrichment of major taxonomic groups relative to source samples (**Fig. 2B and 3A-B**). Comparison of source samples and co-cultures revealed distinct clustering, with *in vitro* community samples tightly clustered relative to samples of source environments (**Fig. S4G, S5G-I, and Table S11**). These observations suggest that even though differences between *in vitro* communities exist, they are substantially smaller than the variations introduced during community formation from source inocula. Thus, even though inoculum source and host strain shape community diversity, the resulting consortia maintain high-level taxonomic coherence across different host strains and inocula.

### Core species identified across model cyanobacterial communities

Although there is no universally accepted definition of a core microbiome, the consistent taxonomic composition and developmental patterns in our *in vitro* cyanobacterial communities (**Fig. 1B, 3A-B, and Fig. S3**) allowed us to define a 25 species core microbiome using genome-resolved data (**See Methods**). These species were reproducibly detected across multiple communities with different cyanobacterial hosts (>= 4 host species) and environmental inocula (>= 2 source environments; **Fig. S6**). Hereinafter we refer to non-core species detected in cultures as auxiliary species and species only detected in environmental samples as source species (**Fig. 2A and Table S9**). Core microbiome species averaged 1.0 ± 1.3% (Range: 0 - 7.3%) of the total community abundance, increasing to 14.7 ± 19.4% (Range: 0 - 82.4%) when considering only non-cyanobacterial taxa (**Fig. 3C**). Individual core species were present in 18.5% - 68.5% of all samples (**Table S9 and Fig. S6**). The majority of core species belonged to taxonomic orders significantly enriched in the L6OTU analysis (**Fig. 2B**) with the most numerous being Rhizobiales (n = 7 species), Rhodobacterales (n = 5 species) and Sphingomonadales (n = 5 species) (**Fig. 3D**). Phylogenetically, core species were closely clustered, with significantly lower phylogenetic distances between their MAGs on average relative to auxiliary or source inoculum species (**Fig. 2A and S7**). Single nucleotide variant (SNV) analysis further confirmed that identical species were independently selected into *in vitro* communities from distinct geographic water sources, with no evidence of strain-sharing between environments (**Fig. S8, Table S13-S14, Supplementary Methods and Results**). Eighteen core species lacked a representative in public databases (**Table S9**), marking an expansion of genomic knowledge for organisms that may occupy a generalized niche in cyanobacterial communities.

**Figure 3.**
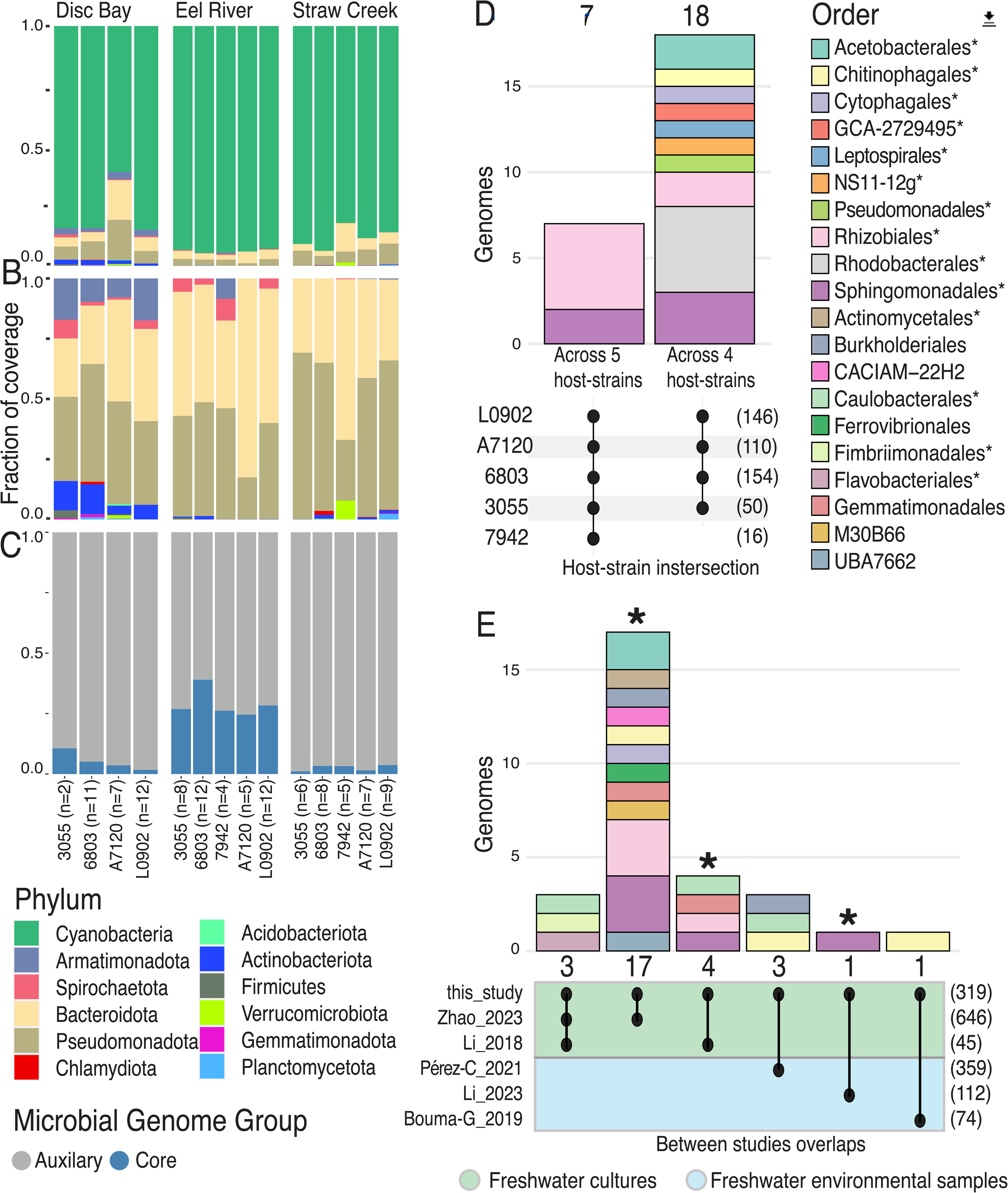
Microbial community composition, core microbiome, and cross community overlap. **(A)** Phylum level relative abundance of *in vitro* communities based on mapping samples to a database of 537 species-level MAGs. Sample compositions are averaged within cyanobacterial strain for each environmental inoculum. **(B)** Phylum level taxonomic composition of *in vitro* communities with cyanobacteria removed. **(C)** Composition of *in vitro* communities with cyanobacteria removed depicting the relative fraction assigned to core and auxiliary microbiome MAGs. **(D)** Cyanobacterial host association and order-level taxonomy of the 25 genomes identified as core microbiome members. The numbers above the colored stacked bars indicate the total number of core species shared. The colored stacked bars represent the order-level taxonomy of these species. Connected dots below each stacked bar indicate the set of cyanobacterial hosts that share the species above. The numbers in parentheses on the right of the plot indicate the total number of species detected in at least three cultures for each host strain. The legend on the right of the figure provides the color code for each order in the stacked bar plot. An asterisk (*) indicates orders that were significantly enriched in co-cultured communities relative to source environment samples (Also See Fig. 2B). **(E)** The number and taxonomy of species shared between the communities of our study and those of other studies where genome-resolved analysis of cyanobacterial consortia was conducted. The numbers above the colored stacked bars indicate the total number species shared. The colored stacked bars indicate the order-level taxonomy of these species. Blue asterisks above the stacked bar plots indicate that a core microbiome species identified in our study was shared. Connected dots below each stacked bar indicate the set of studies that share the species above. Numbers in parentheses on the right of the plot indicate the total number of species recovered in each study. The green and blue color shading under the connected dot plots indicates the type and origin of cyanobacterial community for the compared studies. The coloring of orders in the stacked bar plot of this figure are the same as in Figure 3D.

### Taxonomic similarity between *vitro* and natural phototrophic consortia

We identified 29 (9.1%) shared species, (**Fig. 3E and S9**) between our *in vitro* communities and MAGs compiled from other studies on cyanobacterial consortia (n = 1,843 MAGs). Among these, three MAGs are core microbiome members identified in our study. In total, 22 (76%) shared species also belonged to taxonomic orders enriched during *in vitro* community establishment in this study (**Fig. 2B, 3E**). On average, this study shared more species with other studies (3.2 ± 6.3 species) than the studies did with each other (0.7 ± 2.9 species). Most pairwise comparisons between studies identified no shared species (42/55 comparisons). The species shared with this study were primarily from freshwater-associated cyanobacterial communities, with no species overlap in saltwater, soda lake, or plant rhizosphere communities.

At higher taxonomic levels we observed broad overlap between studies. Out of the 48 observed orders in this study, 37 (77%) were shared with other studies (**Fig. S9B**), with the orders enriched during *in vitro* community establishment in this study being the most commonly shared (*P* = 0.018; Wilcoxon test, **Fig. S9C**). These findings support that the *in vitro* communities developed here reasonably represent natural cyanobacterial consortia and share considerable taxonomic overlap with other phototroph-heterotroph systems.

### Enriched metabolic functions in the core cyanobacterial microbiome

We identified metabolic functions enriched in core microbiome MAGs (n = 25 MAGs) relative to all non-core MAGs (n = 512 MAGs), and a constrained set of non-core MAGs from three phyla in the core microbiome Bacteroidota, Pseudomonadota, and Spirocheaotota (n = 366 MAGs). In both analyses, we found significant enrichment of 1073 and 1065 KEGG orthology groups (KOs), respectively (FDR ≤ 0.05; Fisher test; **Tables S17-S19**). Enriched functional categories included cofactor metabolism (Fisher test; OR_All_ = 1.6, FDR_All_ = 0.007; OR_Constrained_ = 1.9, FDR_Constrained_ = 0.0002) and membrane transport (Fisher test; OR_All_ = 1.7, FDR_All_ = 1.9 e-6; OR_Constrained_ = 1.7, FDR_Constrained_ = 1.1e^-5^). (**Fig. 4A, Table S19**). The constrained analysis mirrored the full analysis and additionally identified signal transduction as (Fisher test; OR_Constrained_ = 1.7, FDR_Constrained_ = 0.006) a significantly overrepresented functional category in the core microbiome (**Fig. 4A**).

**Figure 4.**
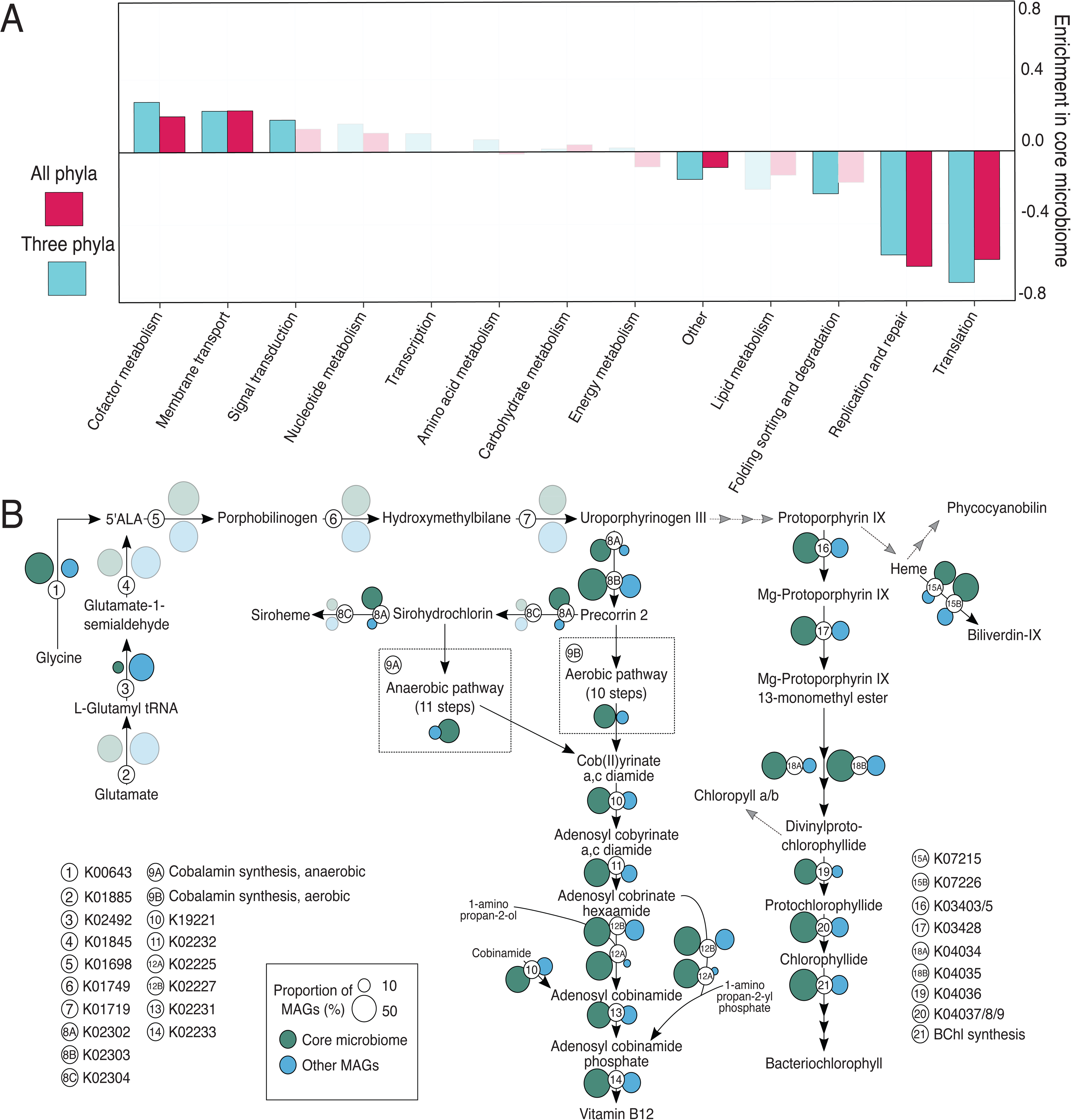
Metabolic functions enriched in the core microbiome. **(A)** Functional categories of statistically over or under enriched KEGG functions in the core microbiome. The enrichment odds-ratio for each functional category is shown for the set of KEGG functions with differential enrichment when the core microbiome MAGs were compared to all 512 non-core MAGs (red) and the 366 non-core MAGs in the phyla *Bacteroidota*, *Proteobacteria*, and *Spirocheaotota* (blue). Bars are solid colors for categories with statistically significant over or under representation (FDR ≤ 0.05; Fisher test) and transparent if not significant. **(B)** Metabolic reaction map for porphyrin compound biosynthetic pathways. Numbers indicate the KEGG KO associated with a reaction. This map shows enrichment data from the comparison of core MAGs to all 512 non-core MAGs. The size of colored circles indicates the percentage of MAGs in core (green; n = 25) or non-core (blue; n = 512) sets that encode the respective reaction. Circles are solid colors if a KEGG KO was detected as significantly over or under enriched in the core microbiome (FDR ≤ 0.05; Fisher test), and transparent if not significant. Also see Tables S16-S18.

Within cofactor metabolism, 39 KOs in the porphyrin metabolic pathway were significantly enriched in core microbiome MAGs including those involved in the production of vitamin B12, chlorophylls, and hemes. The gene encoding 5-aminolevulinate synthase (K00643) which facilitates one-step synthesis of 5-Aminolevulinate (ALA) from glycine was highly enriched in core MAGs (OR_All_ = 9.9, FDR_All_ *=* 6.5 e-6; Fisher test), whereas non-core MAGs favored the more complex three-step pathway from glutamate (K02492; Fisher test; OR_All_ = 24.6, FDR_All_ *=* 3.3 e-8). The gene *cobC* (K02225), involved in vitamin B12 synthesis, showed the strongest core microbiome association (Fisher test; OR_All_ = 24.6, FDR_All_ *=* 3.3 e-8), alongside other enzymes in the vitamin B12 and bacteriochlorophyll synthesis pathways (**Fig. 4B and Table S18**).

We also found enrichment of KOs related to anoxygenic photosystems, with 80% (20/25) of core microbiome members encoding these components (K08929/K08928/K13991; **Fig. 5, Table S18 and S20**). This may indicate that light driven energy production is an important metabolic strategy of cyanobacteria-associated bacteria. Electron donors for these anoxygenic phototrophs are likely organic substrates [91], which agrees with functional predictions. Only one member of the core microbiome, a *Georhizobium* species, encoded RuBisCO for carbon fixation.

**Figure 5.**
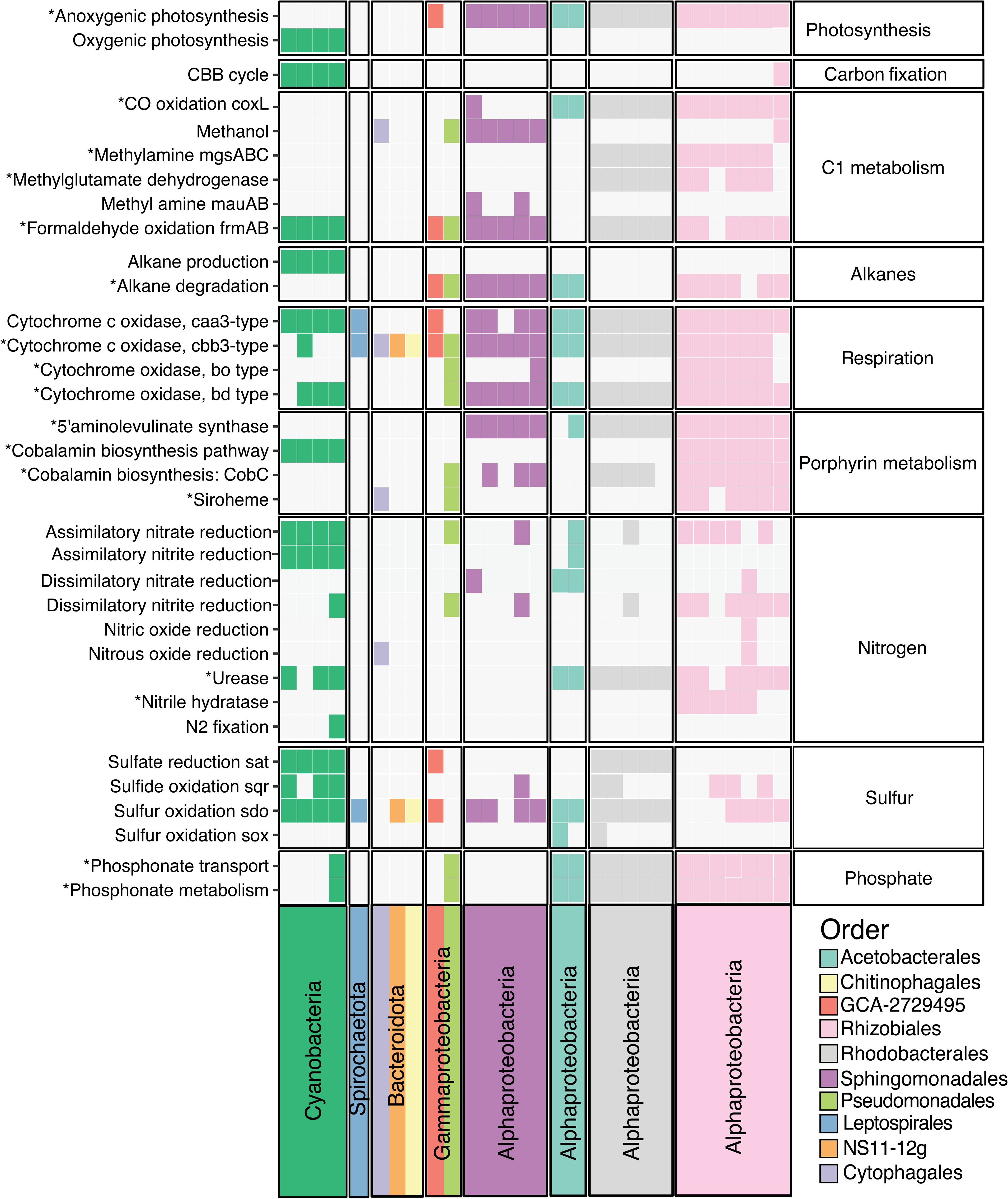
Comparison of metabolic functions in cyanobacterial hosts and core microbiome species. Encoded metabolic potential for specific biogeochemical turnover reactions identified in cyanobacterial host species MAGs and the 25 core microbiome MAGs. Specific metabolic functions (left) are separated by higher level metabolic categories (right). Functions marked with and asterisk (*) had at least one associated KO with significant enrichment in the core microbiome relative to all 512 non-core MAGs. Each column represents an individual MAG in the core microbiome, MAGs are grouped at different phylum levels, and colored by order level taxonomy. Identification of the potential for a specific function in each MAG is indicated by a colored box, and boxes share order and color. Criteria for identifying specific functions can be found in Table S19.

Genes for phosphonate degradation (K06162-6) and transport (K02041-4) were significantly enriched in core MAGs, offering an alternate source of phosphorus to the community. KOs involved in methylamine (K15228-9), nitrile (K01721/K20807), and urea (K01428-30) utilization were also enriched in core MAGs, putatively expanding available carbon and nitrogen sources (**Fig. 5 and Table S18 and S20**). Additionally, many core microbiome MAGs that encoded methylamine to formaldehyde degradation also had genes to oxidize formaldehyde to formate and CO_2_. Finally, core microbiome members were enriched in several genes related to degradation of cyanobacteria-derived long-chain alkanes. All cyanobacterial strains in our study encoded KOs (K14330 and K14331) responsible for long-chain alkane biosynthesis [92, 93], wheres core microbiome members were enriched in genes (*LadA*, *AhyA*, *AlkB*, and *Cyp153*) for the degradation of these compounds [85] (**Fig. 5 and Table S18 and S20**). This suggests that core microbiome species may be utilizing these cyanobacteria-produced alkanes as an energy source.

### Core microbiome MAGs are enriched in putative plasmid elements

Extrachromosomal replicons and plasmids have been shown to encode functions that mediate phototroph-heterotroph symbioses [48]. Here we assessed if there were differences in the frequency and distribution of putative plasmid elements associated with the MAGs resolved in our study. We identified 850 putative plasmid contigs across 537 species-representative MAGs (**Table S21**). Even though 98% of putative plasmids were not circularized, we found 15% encoded plasmid-specific replication domains and 26% encoded proteins for plasmid mobilization (**Fig. S10A**). In our comparison with the comprehensive IMG/PR database, 81 plasmids from our dataset matched entries in the database. Of these 81 matching plasmids, 80% were recovered from metagenomic datasets, primarily from aquatic environments, with only 7 being assessed as putatively complete (**Table S22**).

We found that core microbiome MAGs had a significantly higher proportion of putative plasmid contigs than auxiliary (Dunn Test = 3.7, *P* = 0.0003) or source MAGs (Dunn Test = 6.5, *P* = 0) (**Fig 6A**). As bacteria from the order Rhizobiales are known to encode large numbers of extrachromosomal plasmids [94, 95] we also performed the comparison only between MAGs of this order. We found that core Rhizobiales MAGs contained a significantly larger proportion of putative plasmids than auxiliary (Dunn Test = 3.0, *P* = 0.0036) or source (Dunn Test = 2.4, *P* = 0.0253) MAGs **(Fig. S10D)**.

**Figure 6.**
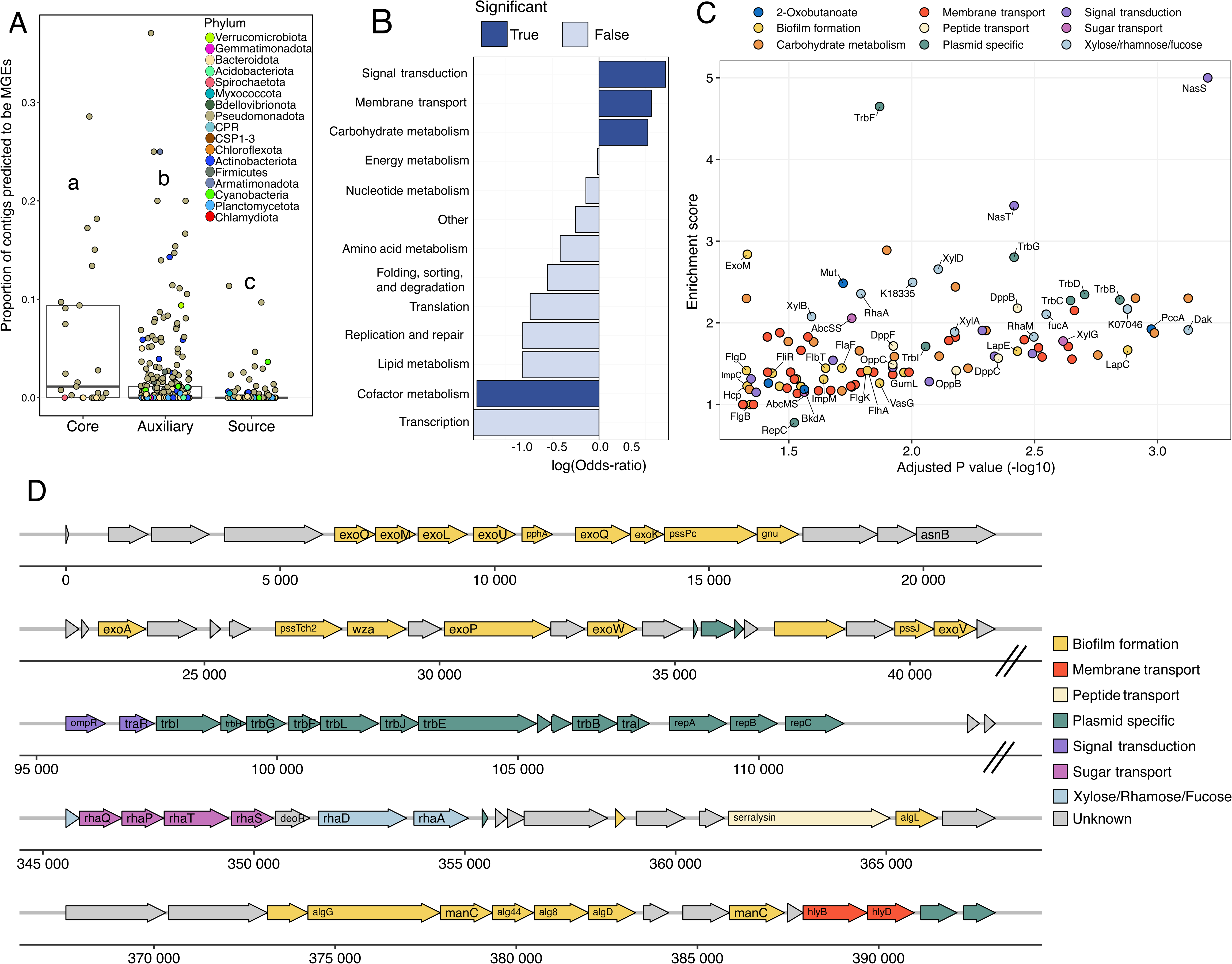
Predicted mobile genetic element distribution and functional potential. **(A)** Proportion of contigs predicted to be mobile genetic elements in each MAG of the core, auxiliary, and source microbiome. Points are colored by phylum level taxonomy of the MAG. Box plots behind points indicate median, first, and third quartiles of data. Statistical differences between groups are indicated by letters, and groups not sharing the same letter are significantly different (Statistic_Core-v-Aux_ = 3.7, FDR < 0.001; Statistic_Core-v-Source_ = 6.5, FDR < 0.0001; Statistic_Aux-v-Source_ = 6.6, FDR < 0.0001; Dunn Test). **(B)** Functional category analysis for the set of KEGG KOs that becomes non-significant when all MGE contigs are removed from MAGs. Functional categories that are significantly over or underrepresented in this set of KOs are colored accordingly (FDR ≤ 0.05; Fisher test). X-axis indicates the natural logarithm of the Fisher test enrichment score. Also see Tables S15, S16 and S20. **(C)** Individual KEGG KOs from membrane transport, carbohydrate metabolism, and signal transduction categories that become non-significant when the enrichment analysis for core microbiome functions is repeated with predicted MGE contigs removed from all MAGs. The x-axis indicates the -log10(FDR) for each KO in the original enrichment analysis. The y-axis indicates the fold change in the odds-ratio for a KO between the original analysis and the analysis conducted with all MGE contigs removed. **(D)** A representative plasmid contig (407,520 bp) co-binned with an *Allorhizobium* MAG from the core microbiome (StrawCreek_S_L0902_W_A_idba_concoct_4). Genes are colored by functional category. Dashed lines indicate a break in the contig for visualization purposes. Also see Table S23.

### Putative plasmids of the core microbiome encode functions facilitating symbioses

To assess functional contribution of putative plasmids in core microbiome MAGs, we repeated our functional enrichment analysis, excluding plasmid-predicted contigs from all MAGs. We identified 197 KEGG Orthology (KO) groups that were no longer significantly enriched in the core microbiome (**Tables S18 and S19**), indicating that these KOs were enriched on putative plasmids. Functional category analysis of this KO set revealed significant enrichment of signal transduction (Fisher test; OR = 0.7, FDR *=* 0.038), membrane transport (Fisher test; OR = 0.6, FDR *=* 0.038), and carbohydrate metabolism (Fisher test; OR = 0.5, FDR *=* 0.049) on putative plasmids of the core microbiome (**Fig. 6B**).

Enriched plasmid-specific functions on core microbiome plasmids (**Fig. 6C and Table S18**) included those for plasmid replication (*repC*) and conjugal transfer (*TrbBCDFGI*). Carbohydrate metabolic functions included those mediating the interconversion and metabolism of sugars commonly found in cyanobacterial exopolysaccharides [96, 97], such as xylose (*XylABD*), rhamnose (*RhaAM*), and fucose (K18335, K07046, and *FucA*). Transporters for importing xylose (*XylG*) and simple sugars (*AbcSS* and *AbcMS*) were also enriched. Additionally, putative plasmids in the core microbiome may enable the utilization of peptides as alternative carbon and nitrogen sources as evidenced by the enrichment of dipeptide (*DppBCF*) and oligopeptide (*OppBC*) transport systems, and genes (*Mut*, *PccA*, and *BkdAB*) involved in converting 2-oxobutanoate to succinyl-CoA, linking amino acid degradation to the TCA cycle.

We identified plasmid-enriched genes that may mediate symbiotic interactions and biofilm formation. These genes include multiple flagellar structural proteins (FlgBDK and FlhAR), a flagellar regulatory protein (RpoD), and components of an Imp-like type VI secretion system (ImpBCFM) (**Fig. 6C and Table S18**). Furthermore, we identified a set of plasmid-enriched KOs involved in exopolysaccharide biosynthesis (*exoM* and *GumL*). We discovered comprehensive gene clusters for the biosynthesis of succinoglycan and alginate on a large (407 kb) putative plasmid (**Fig. 6D and Table S23**) associated with a Rhizobiales MAG from the core microbiome (StrawCreek_S_L0902_W_A_idba_concoct_4). Although this element was not fully circularized, it encoded plasmid-specific replication factors (*repABC*) and a complete conjugal transfer system, including a relaxase protein (TraI) and type IV secretion apparatus (*trb*).

## Discussion

In this work, we built on existing approaches [21–23, 25, 26, 36–38] to generate and analyze 108 *in vitro* cyanobacterial consortia using well-characterized, model cyanobacterial hosts. This approach provides three core advances for the *in vitro* study of cyanobacterial communities.

First, leveraging cyanobacterial host species that serve as key model systems in cyanobacterial research [43] significantly enhances our previously poor understanding of the microbiomes they recruit. Second, extensive replication enabled the identification of a core microbiome across multiple taxonomically diverse cyanobacterial hosts. Third, leveraging this core microbiome, we confirmed, statistically rather than solely by observation, enrichment of functional traits associated with core species, and, furthermore, uncovered metabolic functions and the potential for plasmids to mediate phototroph-heterotroph symbioses in these systems.

The cyanobacterial hosts used in this study were selected for their diversity and importance as model organisms for advancing our understanding of cyanobacterial biology [41, 42, 98–103]. Despite their value as models, little is known about their behavior in ecological contexts.

Whereas previous studies have utilized some of these species in two-member co-culture systems [8, 32, 33, 35], this study provides the insights into more complex microbial communities potentially recruited by these species from natural environments. We provide strong evidence that not only can *in vitro* communities rapidly stabilize around these model hosts (**Fig. 1B-D and S3**), but that these communities exhibit strong taxonomic overlaps with freshwater cyanobacteria in natural habitats [28, 30], cultivated in lower-throughput laboratory systems [23, 78], and share higher level taxonomic overlap with plant rhizosphere associated communities (**Fig. 3E, S9**). Furthermore, evidence supporting host-specific selective pressures was observed, as species co-occurring with the same host exhibited significantly higher strain-level ANI across co-cultures compared to the same species associated with different hosts **(Fig. S9)**.

By inducing co-culture diversity through variation of both cyanobacterial host species and environmental inoculum sources, consistently shared features could be robustly identified. This diversity was leveraged across consortia to define a 25-species core microbiome (**Fig. 3D**) identified across five model cyanobacterial host strains. Despite the limitation of using freshwater cyanobacterial species exclusively, the core microbiome identified here provides a foundation for understanding organisms that commonly co-occur across an array of taxonomically and functionally diverse cyanobacteria. Although some core species identified here were also present in cyanobacterial consortia from other studies (**Fig. 3E**), our hypothesis is that the specific identities of these core microbiome members are less critical than the functional traits and “microbial archetypes” they represent. Therefore, comparing the genomic content of core and non-core species provides a framework for statistically identifying functional traits that play a broader role in supporting cyanobacterial ecosystems.

Functional genomic analysis of the core microbiome revealed a significant enrichment of metabolic pathways associated with the biosynthesis and potential cross-feeding of resource-intensive compounds (e.g., vitamin B12), the enhancement of nitrogen and phosphorus availability, and anoxygenic phototrophy (**Figs. 4 and 5**). Even though some of these pathways, such as anoxygenic photosynthesis [28, 30], have been previously reported [12, 16] this study provides robust statistical evidence for their specific enrichment within a core microbiome.

Collectively, these findings support the hypothesis that core microbiome functions facilitate metabolic cooperation centered around phototrophy. This is further reinforced by the predominance of porphyrin cofactor metabolism among enriched functions, including numerous enzymes involved in vitamin B12 and bacteriochlorophyll biosynthesis pathways (**Fig. 4**). The core microbiome exhibits enrichment in ALA synthase, which enables a more energetically efficient route for ALA production from glycine, potentially advantageous in communities where porphyrin compounds are critical resources. The importance of such exchanges is underscored by studies demonstrating enhanced cyanobacterial growth when supplemented with the porphyrin precursor ALA [104]. Although vitamin B12 cross-feeding in microbial communities has been studied intensively [105–107], the shared production and exchange of other energetically demanding and metabolically linked molecules, such as chlorophyll precursors, remain poorly characterized. The enrichment of anoxygenic phototrophy within the core microbiome, consistent with observations from other studies [91], may suggest that non-competitive photosynthetic [108] partnerships are important in cyanobacterial microbiomes for exchanging essential and costly metabolic precursors common to all organisms. Finally, although multiple studies have posited the function of cyanobacterial long-chain alkanes [109], this study shows the enrichment of long-chain alkane degradation pathways in core microbiome organisms (**Fig. 5**). This capability was specifically prevalent in those core microbiome organisms that also encoded phototropic capabilities. This observation would support that cyanobacteria produce these compounds to specifically select and modulate their surrounding communities by creating a metabolic niche.

Even though plasmids have been studied in microbial symbioses [110, 111], their roles within cyanobacterial consortia remain largely unexplored. Here, core microbiome genomes exhibited a higher prevalence of plasmids compared to non-core members (**Fig. 6**). Functionally core microbiome plasmids encoded gene clusters linked to the biosynthesis of exopolysaccharides such as succinoglycan, alginate, and xanthan gum-like compounds, polymers known to play essential roles in plant-bacterial interactions [110, 112]. Succinoglycan is critical for nitrogen-fixing Rhizobiales [113, 114], and alginate forms hydrogels that enhance biofilm formation [115–117]. In particular, alginate encapsulation of *Synechococcus elongatus* has been shown to improve both sucrose export and co-culture stability in laboratory settings [32]. Additionally, core microbiome plasmids contained a two-component nitrate regulatory system (*NasST*), gene clusters for mobility machinery, and Type VI secretion systems, features commonly found in plant-associated bacteria [118], suggesting potential roles in colonization, competition, and biofilm formation. Beyond colonization and biofilm forming functionality, plasmids in core microbiome species encoded metabolic pathways for pentose and hexose sugar utilization, including rhamnose and xylose metabolism, key components of cyanobacterial exopolysaccharides [96, 97] (**Fig. 6, S10, Tables S17-18, S22**). Collectively, these findings support the hypothesis that plasmids contribute to core microbiome establishment and maintenance by enhancing biofilm formation and niche adaptation.The data further highlight adaptive advantages plasmids may confer within cyanobacterial consortia, providing new insights into how plasmids may shape these microbial interactions.

Even though compelling, several limitations of our approach warrant discussion. The absence of cyanobacteria-free controls, limited our ability to disentangle effects solely attributable to media or laboratory conditions. However, we observed substantial taxonomic overlap (**Fig. 3E and S9**) between our co-cultures and other freshwater cyanobacterial and rhizosphere consortia [6, 28, 45, 46, 103]. Furthermore, the use of naturally sourced inocula carried the potential for introducing diverse eukaryotic organisms, including flagellates, ciliates, fungi, and phototrophic diatoms and algae. Although we rigorously screened cultures for readily detectable eukaryotes (**See Methods**), we did not assess the diversity or function of eukaryotic species present within final co-cultures, which on average constituted 0.64±1.89 % of assembled data on an assembled contig size basis (**Table S2**). Our objective was to focus on the prokaryotic interactions of model cyanobacteria, given the limited existing data on these hosts’ recruited microbiomes. Despite these constraints, our findings and related work [19] underscore the value of *in vitro* cyanobacterial model systems for revealing insights into phototroph–heterotroph interactions. Although methods for cultivating phytoplankton consortia are well-established [19, 23, 28, 38, 47], our framework and biobanked stable co-cultures offer a controlled yet ecologically relevant platform to advance understanding of freshwater microbial community dynamics. Future investigations using complementary methods such as long-read sequencing, mesocosm experiments, cyanobacteria-free controls, and inclusion of eukaryotic organisms, will help refine our understanding of aquatic microbiome functionality.

## Supporting information

Supplementary Materials Methods and Results

## Data Availability

Raw sequencing read data generated as part of this project are available under the NCBI BioProject accession number PRJNA1271998. All metagenome bins and putative plasmid sequences recovered in this study have been deposited in FigShare under the following DOIs: 10.6084/m9.figshare.26148823 and 10.6084/m9.figshare.26148811

## Code Availability

Code used in the analysis for this paper are available at the following GitHub repository: https://github.com/SDmetagenomics/Kust_Cyano_2023

## Reporting Summary

Further information on research design is available in the Nature Research Reporting Summary linked to this article.

## Acknowledgements

We would like to thank Prof. Susan Golden, Prof. James Golden, and Dr. Arnaud Taton for providing the cyanobacterial strains used in this study. Eel River sampling occurred at the UC Angelo Coast Range Reserve. Funding for this project was provided in part by the Gordon and Betty Moore Foundation (GBMF9321). Funding for this project was also provided in part by the Shurl and Kay Curci Foundation.

## Author Contributions

AK, JFB, and SD conceived overall experimental concepts and design. AK and SD designed the sampling and cultivation strategy. AK, KBG, and SD collected water samples. AK, CCP, WL, and SD performed cultivation and sampling. AK, CCP, NK, and SD performed sequencing and sequence processing. AK and SD performed sequence assembly and genomic binning. AK, JZ, and SD performed annotation and statistical analysis on amplicon and metagenomic datasets. AK, JZ, and SD wrote the manuscript. AK, JZ, JFB, and SD edited the manuscript

## Competing Interests

The authors declare no competing interests.

## Corresponding Author

All correspondence should be addressed to Spencer Diamond

## Notes

https://www.ncbi.nlm.nih.gov/bioproject?term=PRJNA1271998

https://doi.org/10.6084/m9.figshare.26148823

https://doi.org/10.6084/m9.figshare.26148811

https://github.com/SDmetagenomics/Kust_Cyano_2023

### Competing Interest Statement

The authors have declared no competing interest.

### Summary of Updates

In our revision, we have significantly revised the work to: (i) better address novelty of the approach in the context of prior work, (ii) address technical concerns regarding sequencing, (iii) add considerable detail with respect to culture contamination monitoring and study inclusion, (iv) included new analyses to compare our systems with other freshwater cyanobacterial systems evaluated at genome-level resolution.

