## Supplementary Materials Methods and Results for "Model cyanobacterial consortia reveal a consistent core microbiome independent of inoculation source or cyanobacterial host species"

### Supplementary Methods

#### Selection of model cyanobacterial strains and associated species level genomes

*Synechococcus elongatus* UTEX 3055 (S3055) is a unicellular strain that forms biofilms and is capable of phototactic motility with a unique bidirectional photoreceptor [1]. This is a strain variant of *Synecococcus elongatus* PCC 7942 which does not strongly exhibit these capabilities

*Synechococcus elongatus* PCC 7942 (S7942) is a more widely studied unicellular strain variant of *Synechococcus elongatus* UTEX 3055 that exhibits a self-suppression mechanism that allows for constitutive planktonic growth under standard laboratory conditions [2]. This strain has been investigated for its unique characteristics, such as the circadian clock mechanism [3] and gene expression patterns under different environmental conditions. Specifically, it has been engineered for glucose utilization, CO2 fixation, and chemical production [4], production of valuable compounds like α-farnesene through metabolic engineering [5], development of synthetic microbial consortia for biotransformation processes [6], and mixotrophy under natural light conditions for improved feedstock production [7].

*Synechocystis* spp., PCC 6803 (S6803) is a unicellular strain, the first cyanobacterial strain that had it’s whole genome sequenced [6, 8], and is one of the most extensively studied cyanobacteria. It has been utilized in various biotechnological applications and synthetic biology studies, such as biofuel production and metabolic engineering. Specifically, it has been studied for its response to osmotic shock and the role of the Na+-dependent K+ uptake Ktr system in cell adaptation, synthesis of polyhydroxyalkanoate (PHA) and ethanol, and stress acclimation [9–13].

*Nostoc/Anabaena* spp., PCC 7120 (A7120) is a filamentous cyanobacterium with the capability to differentiate specialized nitrogen-fixing cells known as heterocysts [14]. This strain is a model for cellular differentiation, nitrogen fixation, and hydrogen production [15, 16]. It has been used in various biotechnological applications such as oxidative stress tolerance [17, 18], photobiological hydrogen production [19], response to environmental stressors [20], and enhanced photosynthesis.

*Leptolyngbya* spp., BL0902 (L0902) is a robust filamentous cyanobacterial strain known for its high biomass production capabilities and tolerance to a wide range of salinities and light intensities, making it a promising candidate for environmental and biotechnological applications [21]. Its rapid growth rates, the ability to form homogeneous suspensions in liquid media, which facilitates the use of optical density (OD) measurements for growth rate estimation, environmental resilience, and genetic tractability is valuable for controlled model synthetic community study [21, 22].

MAGs that were used as species-level representatives for each strain are as follows (**Table S9**):

- L0902 = unicom2021_Dbay_Mouth_L0902_BW_C_idba_metabat.8
- S7942/3055 = unicom2021_Eel_Fox_3055_W_B_spades_maxbin.002
- S6803 = unicom2021_StrawCreek_S_6803_BW_A_idba_maxbin.001
- A7120 = unicom2021_StrawCreek_S_A7120_W_A_spades_metabat.3

**Experimental setup, inoculation, and preservation of cyanobacterial co-cultures**

Cyanobacterial strains were initially grown axenically in 250 mL Erlenmeyer flasks with 100 mL of BG-11 medium [23] (supplemented by nitrate, but not vitamins) at 24°C in a 12:12 light:dark cycle. Strain axcenicity prior to the experiment was confirmed by fluorescence microscopy after DAPI staining and by incubation of samples taken from cultures on LB agar plates in the dark at 30°C.

Water samples were collected from three freshwater bodies located in Discovery Bay, California ("Discovery Bay"), Branscomb, California ("Eel River"), and Berkeley, California ("Strawberry Creek"). These environments were selected due to their distinct sets of abiotic parameters (**Fig. 1A and Table S1**) that still fall within acceptable ranges for each cyanobacterial species used as a host for *in vitro* community cultivation. Abiotic parameters, including pH, temperature, and nutrient concentrations, were measured (**Table S1**). Metagenomic analysis of the microbial communities from each site confirmed that each freshwater body harbored distinct microbial compositions dominated by common freshwater bacterial phyla, such as Pseudomonadota, Bacteroidota, and Verrucomicrobiota (**Fig. S2**) [24, 25] . These distinct microbial communities provided varied inocula for the cyanobacterial strains. Bacterial fractions from six surface freshwater locations (two sites per location) (**Fig 1A**, **Table S1**) were independently concentrated using a multi-step filtration method. Water was pre-filtered through 20 and 5 µm filters to discard fractions bigger than 5 µm to avoid bigger freshwater organisms. The filtrate was then passed through 0.2 and 0.1 µm polyethersulfone filters until saturation was reached, indicating sufficient bacterial biomass had been collected. Approximately 2 L of water was required to saturate filters from all three locations (n = 6 sites), indicating sufficient bacterial biomass had been collected. The bacterial biomass on the 0.1 µm and 0.2 µm filters was used for DNA extraction and inoculation of axenic cyanobacterial cultures. Filters were bisected, one half from each filter size (0.2 and 0.1 µm) was directly stored at -80°C for DNA extraction. The remaining halves were then transferred into 30 mL of BG11 medium. These filters underwent a 30-minute vortexing at maximum speed to ensure the dissociation of bacterial cells from the filter. The resulting bacterial suspensions from the 0.1 µm and 0.2 µm filters were combined, and the optical density 600 nm ($OD_{600}$) was measured. Volume of bacterial suspension was then added to freshly inoculated axenic cyanobacterial strains ($OD_{680}$=0.2) to achieve a normalized bacterial inoculation density of $OD_{600}$ = 0.01. This filtering and inoculation process was repeated three times for each of the six sites where water was collected, resulting in three biological inoculation replicates for each condition we evaluated (A, B, C). This generated a total of 180 initial consortia (n = 5 cyanobacterial hosts, n = 6 sub-locations, n = 2 passaging intervals, n = 3 replicates).

Following inoculation, the *in vitro* cyanobacterial communities were grown at 24°C in a 12:12 light:dark cycle and passaged into fresh BG-11 medium at either weekly or bi-weekly intervals. At the start and end of each passage interval, samples were collected to determine the optical density of the co-culture at 680, 700, and 750 nm. A volume of each co-culture from the previous passage was inoculated into fresh BG-11 such that $OD_{680}$ = 0.2. Additionally at each passage interval biomass was collected for DNA extraction (2 × 2 mL), and fixation for microscopy (1 mL). Biomass for DNA extraction was centrifuged for 20 min at 11,000xg at room temperature, frozen, and stored at -80°C until DNA extraction.

At the final time point, biomass from 120 cultures that were retained until the day 84 was cryopreserved in 4% DMSO [26]. These preserved cultures were used to evaluate viable regrowth and community stability, facilitating the development of stable cyanobacterial consortia capable of long-term storage and future reconstitution.

**Assessment of cultures contamination and retention**

To ensure culture integrity and minimize contamination, we implemented a multi-tiered assessment approach throughout the passaging process. This included visual inspection, microscopic evaluation, metagenomic analysis of host dominance, and a newly introduced estimation of the eukaryotic fraction. Below, we outline each of these approaches in detail.

(i) Visual assessment: At each passaging time point, all cultures were visually assessed for discoloration (e.g., yellowing), low turbidity/limited growth, and the presence of unusual debris. Cultures exhibiting any of these characteristics were subjected to further microscopic evaluation to determine potential contamination or other abnormalities.

(ii) Microscopic assessment: Cultures displaying visual abnormalities, as well as a randomly selected subset of 20 additional cultures per passaging interval, were examined using light microscopy. Non-host cyanobacterial eukaryotic contaminants, including algae, diatoms, and protist-like organisms, were readily detectable through this method. Any culture in which these contaminants were observed in any quantity was removed from the study. For instance, a culture containing >50% eukaryotic algae would be discontinued. Additionally, cultures harboring cyanobacteria with obvious morphologies distinct from the expected host were also excluded. If no contaminating organisms were detected and only cyanobacteria matching the expected host morphology were present, the culture was retained for an additional passage and re-evaluated at the subsequent passaging interval.

In cases where discoloration or unexpected behavior was observed without an immediately identifiable cause, cultures were given an additional passage in fresh media. If they failed to recover, they were discontinued. A qualitative assessment of the 72 cultures that ultimately failed revealed that most contained protozoan contaminants, as identified via light microscopy, suggesting predation as a potential cause of culture decline—an established challenge in dense cyanobacterial cultures exposed to environmental microbes [27].

(iii) Metagenomic host dominance assessment: For cultures that were retained until the 84-day time point, we evaluated the fractional dominance and identity of the present cyanobacterial species. Cultures containing mixed cyanobacterial species that did not match the expected host were removed from the experiment. Specifically, any sample where the anticipated host cyanobacterium comprised <90% of the total cyanobacterial fraction was discontinued. This led to the removal of 12 cultures, with the detailed results of this assessment provided in Supplement **Fig. S2.**

(iv) Estimation of eukaryotic fraction: To assess the potential presence of eukaryotic organisms in our retained co-cultures, we performed an additional analysis on assembled shotgun metagenomic contigs. We employed Whokaryote+Tiara [28], a classifier designed to minimize taxonomic bias in eukaryotic identification. To ensure high specificity, we applied a stringent filtering threshold, retaining only contigs ≥ 20 Kbp for classification. We then quantified the proportion of contigs classified as eukaryotic in assemblies on a length basis (e.g. the percentage of assembly length that was classified as eukaryotic), providing an estimation of the eukaryotic fraction within each metagenome. This analysis revealed that only three cultures contained more than 5% eukaryotic contamination, with an average eukaryotic fraction of 0.64±1.89% across all 108 co-culture communities retained in the experiment. A summary of these findings is provided in **Table S2.**

**DNA extraction for shotgun metagenomics and sequencing**

DNA for shotgun metagenomic sequencing was extracted from the frozen combined 0.2 and 0.1 µm polyethersulfone filter set with the biomass from each environmental sub-location (n = 6 samples), the bacterial inocula biomass for each sub-location directly used to inoculate axenic cyanobacterial cultures (n = 6 samples), the frozen 5 µm polyethersulfone filter biomass from both Discovery Bay sub-locations (n = 2 samples), and the frozen biomass of all co-culture at day 84 of the experiment (n = 120 samples). DNA extraction was performed with the Qiagen DNeasy PowerSoil Pro Kit (Qiagen, USA) with modification. Briefly, an additional initial lysis step was performed where samples were first heated in CD1 solution at 65°C for 30 minutes prior to the subsequent bead beating step. The DNA concentrations of the samples were quantified using a Quibit fluorometer (Thermo Fisher Scientific). DNA extracted from co-cultures (n = 120 samples) and from source material samples (filters and bacterial inocula; n = 14 samples) were subsequently sent for sequencing using two different sequencing strategies. DNA from source material samples was sent to the QB3 Genomics Laboratory at the University of California, Berkeley for library preparation and paired-end sequencing on a NovaSeq System (Illumina) using a 250 bp X 2 read length at an approximate depth of 10 Gbp per sample. Raw DNA from co-culture samples were sent to Novogene (Guangzhou, China) for library preparation and paired-end sequencing on a NovaSeq System (Illumina) using a 150 bp X 2 read length at an approximate depth of 10 Gbp per sample. We used longer 250 bp reads for environmental samples, as they were expected to be more complex and diverse than co-cultures, allowing for better resolution of ambiguity during assembly [29], the more cost-effective 150 bp reads were used for co-culture samples.

**DNA extraction for 16S rRNA gene amplicon sequencing**

Genomic DNA was extracted from the biomass of cyanobacterial cultures at each passage using the method as previously described by Clerico et al. (2007). A total of 1,941 samples were sequenced in 10 batches. Samples were amplified for the V4–V5 hyper-variable regions of the 16S rRNA gene using primers 515F (5′-GCTCTTCCGATCTGTGYCAGCMGCCGCGGTAA-3’) and 926R (5’-GCTCTTCCGATCTCCGYCAATTYMTTTRAGTTT-3’), both containing Illumina adapter sequences. Each PCR reaction included 10 ng of template DNA, 0.5 ng of the ZymoBIOMICS® Microbial Community Standard (ZymoResearch), and the Phusion® High-Fidelity PCR Master Mix (Thermo Fisher Scientific).

To assist with the identification of standards in 16S rRNA gene amplicon data, we constructed and included a dilution curve of the standard in each amplicon sequencing batch. Briefly, DNA extracted from a single cyanobacterial community sample was randomly selected from each sequencing batch and was spiked with varying concentrations of the ZymoBIOMICS® Microbial Community Standard (2, 1, 0.5, 0.25, 0.1 ng). This spike-in was prepared in duplicate resulting in 10 spike-in samples per sequencing batch and a total of 90 spike-in samples across all sequencing runs. Additionally, 1 sample per run included only 0.5 ng of the standard without any cyanobacterial community DNA, and 3 wells contained only the PCR reaction mix without any template.

The PCR products were purified using SPRI AMPure XP beads (Beckman Coulter) to select for a fragment size of approximately 400 bp. The purified PCR products were then quantified using the Quant-iT High-Sensitivity dsDNA Assay Kit (Thermo Fisher Scientific) in a Tecan Spark® Multimode Microplate reader (Tecan). Following purification and quantification, the PCR products were sent to the Innovative Genomic Institute’s high-throughput sequencing core. Here, they were sequenced on an MiSeq System (Illumina) using a read length of 2 x 300 bp and an expected read depth of 50,000 reads per sample. The sequencing depth was chosen to sufficiently capture the data necessary for frequent monitoring of community changes, complementing whole genome sequencing, which was employed for more in-depth analysis.

**16S rRNA gene data processing and downstream statistical analysis**

16S rRNA gene amplicon sequencing was used to track longitudinal changes in community composition, diversity, and stability across all co-culture samples during every passaging step, as high-coverage shotgun metagenomics would have been cost-prohibitive. All samples were used to identify amplicon sequence variants (ASVs) using the USEARCH-UNOISE3 pipeline [30, 31] with modifications. Briefly, we first assessed expected error rates and read orientation of raw reads using the `usearch -fastx_info` and `usearch -search_oligodb` commands with default parameters. Forward and reverse reads from each sample were then merged and aggregated using following command: `usearch -fastq_mergepairs *_R1_*.fastq -relabel @ -fastqout merged.fq -fastq_maxdiffs 10 -fastq_pctid 90`. Merged reads were then placed into the same orientation by searching against the SILVA_v138 SSU reference database [32] using the following command: `usearch -orient merged.fq -db SILVA_138_SSURef_NR99_tax_silva_trunc.fasta -fastqout oriented.fq -tabbedout orient.txt`. We subsequently removed reads from the analysis where the forward and reverse V4–V5 primers did not fully and identically match forward and reverse reads respectively by first searching for these primer sequences in our merged reads using the following command: `usearch -search_oligodb merged.fq -db primers_V4-V5.fa -strand both -threads 24 -userout merged_primer_hits.txt -userfields query+target+qlo+qhi+qstrand+diffs+trowdots+pairs`. We subsequently identified and filtered these problematic merged reads from the dataset using a custom R script: `Rscript filter_usearch_oligo_match.R -i merged.fq -H merged_primer_hits.txt`. Primers were then hard trimmed from the merged reads using the following command: `usearch -fastx_truncate merged_filtered.fq -stripleft 19 -stripright 20 -fastqout stripped.fq`. Merged reads with bases called as N or with an expected error rate of > 1 were then removed from the data with the following command: `usearch -fastq_filter stripped.fq -fastq_maxee 1.0 -fastq_maxns 0 -fastaout filtered.fa`. Unique sequences and their total count in the full dataset were then identified using the following command: `usearch -fastx_uniques filtered.fa -fastaout uniques.fa -strand both -sizeout -relabel Uniq -minuniquesize 1 -uc uniques.uc`. We then identified ASVs in our dataset and required an ASV to have at least 100 observed counts across all samples (0.005% of total sequenced reads) using the following command: `usearch -unoise3 uniques.fa -zotus zotus.fa -tabbedout unoise3.txt -minsize 100`. An ASV count table was generated by mapping all sequences following primer removal to ASV sequences using the following command: `usearch -otutab stripped.fq -zotus zotus.fa -otutabout zotutab.txt -mapout zmap.txt`. Our ASV identification pipeline removed 6 samples and generated 2126 ASVs from 1,935 samples. We were able to assign 87% of sequenced amplicons generated in our study to this ASV set. Taxonomy was assigned to ASVs by first training a custom naive-bayes classifier using the SILVA_v138 database [32] truncated to the SSU V4-V5 region using the QIIME package [32, 33]. Subsequently ASVs were taxonomically classified using the trained classifier. Finally, we identified ASVs associated with spike-in standards using a custom R script that performs two steps: (1) The sequence identity of ASVs to a database of 16S rRNA gene sequences from known spike-in organisms is calculated using ublast; (2) Robust regression analysis is applied to evaluate the slope of ASV counts relative to the known mass of spike-in standards across sets of spike-in samples included in each amplicon sequencing run. ASVs with a sequence identity ≥ 90% to known spike-in 16S sequences and a statistically significant positive slope (p-val of regression ≤ 0.05) were identified as standards (**Tables S3-S5)**; See Data Availability, and Code Availability for additional information.

Amplicon samples were subset and filtered for downstream analysis to achieve three criteria: (i) to make amplicon and MAG-resolved analyses congruent; (ii) to allow for direct comparison between covariates of weekly and bi-weekly passaging regimes; (iii) to remove select samples of low quality. Briefly, we first subset amplicon samples to the same set of cultures (e.g. combinations of source environment, cyanobacterial host strain, biological replicate, and passage rate) used in MAG resolved metagenomic analysis (n = 108 individual cultures; n = 1240 amplicon samples; See Below). Genome-resolved metagenomic analysis identified cultures that met visual criteria for retention during passaging but did not satisfy community composition quality control criteria (See Below).

Subsequently, we only retained and analyzed amplicon samples collected across these 108 cultures at bi-weekly intervals (14, 28, 42, 56, 70, 84, 91, 105, and 119 d) to allow for direct comparisons between weekly and bi-weekly passaging regimes (n = 108 individual cultures; n = 888 amplicon samples). Finally, we removed individual amplicon samples with ≤ 1000 total ASV counts (n = 1 amplicon sample) and samples where the most abundant ASV was not the expected host cyanobacterium (n = 18 amplicon samples). Following sample selection and filtering amplicon analysis was performed on a final set of 869 samples (**Table S3 - S6**).

Statistical analysis of amplicon data was conducted in R v4.2.3 (See Code Availability). The relative abundance of order level taxa in cultures was calculated as the fraction of total counts assigned to ASVs grouped at the order taxonomic level per sample after ASVs identified as internal standards were removed. Relative abundance was averaged across all samples displayed in respective plots. Area plots were constructed using the ggplot package [34] in R.

Longitudinal alpha-diversity analysis was conducted on rarefied ASV counts with ASVs identified as internal standards removed. Briefly, ASV matrices were rarefied using the GUniFrac package in R [35] to the depth of the sample with the lowest number of counts (4746 and 642 counts for all ASVs and ASVs with cyanobacteria removed respectively). The mean number of counts per sample including cyanobacterial ASVs was 46,098, and the mean number of counts per sample without cyanobacterial ASVs 14,565. Richness and Shannon diversity metrics were calculated using the microbiome package in R [36]. The impact of inoculation source, cyanobacterial host, passage rate, and cultivation time on Richness and Shannon diversity were evaluated using linear mixed effects models implemented with the lme4 package in R [37]. All models took the following form: `[alpha-diversity metric] ~ General Site + Strain + Passage Rate + Total Days + (1|Tube Number)`. Total cultivation time in days was modeled as a factor rather than a continuous variable to allow for comparisons between time points. Variation for each cyanobacterial culture (e.g. tube number; n = 108 cultures) was included as a random effect to account for the individual variance in each culture over longitudinal sampling. Evaluation of covariate significance in models was performed using Type III Analysis of Variance with the `anova` function in R. Pseudo-R-squared values for models were calculated with the MuMIn package [38, 39] in R. Estimated marginal means for alpha-diversity values at each time point were generated and statistically significant differences between all pairwise marginal means were evaluated using the emmeans package [40] in R.

Compact letter displays for significantly different groups were generated using the multcomp package in R.

Lagged beta-diversity analysis was conducted on additive log ratio (ALR) transformed ASV count data with and without ASVs taxonomically classified as cyanobacteria. Briefly, we first removed ASVs that had positive counts in ≤ 2 samples. We subsequently performed zero imputation on count matrices using the zCompositions package [41] in R. ALR transformation was then applied using a custom R function (See Code Availability) that utilizes the counts of ASVs classified as standards in each sample to calculate the geometric means for each sample. Aitchison distance between samples was then calculated as the Euclidean distance between ALR transformed samples using the vegan package [38] in R. For each individual cyanobacterial culture (n = 108 cultures) we captured the Atchison distance between each sample and its previous time point (e.g. 14 d ← 28 d). This resulted in 8 time point comparisons across the time course for each cyanobacterial culture. The impact of inoculation source, cyanobacterial host, passage rate, and cultivation time on beta-diversity between preceding time points were evaluated using linear mixed effects models implemented with the lme4 package in R [37]. Subsequent statistical analysis was conducted identically to alpha-diversity analysis above with cyanobacterial culture number included as a random effect in the model. Complete statistical outputs from the diversity modeling analysis are available in the supplementary materials **(Table S5**).

Differential abundance of ASVs between the 14 d → 84 d and the 84 d → 119 d time points was evaluated using the Maaslin2 package [37, 42] in R. Briefly, ASVs with < 10 positive counts across all samples were first removed (n = 1557 remaining ASVs). The Maaslin2 function was then run on the resulting count matrix with the following parameters: `min_abundance = 0, min_prevalence = 0, min_variance = 0, analysis_method = "LM", max_significance = 0.05, fixed_effects = c("General_Site", "Strain", "Passage_Rate", "Total_Days_Fac"), reference = c("Total_Days_Fac,84"), random_effects = c("Tube_No"), correction ="BH", cores = 12, plot_heatmap = F, plot_scatter = F`. ASVs with significant differences were required to have a FDR ≤ 0.05. All differential abundance statistics are available in the supplementary materials **(Table S6)**.

Community diversity and taxonomic composition were tracked over time to evaluate stabilization, with minimal change observed after 28 days **(Fig. 1B-D, Fig. S3, and Tables S3-S6).** Diversity metrics and taxonomic compositions were monitored to compare the effects of weekly and bi-weekly passaging regimes, as well as cryopreservation and revival on community stability **(Fig. S3, Table S3-S6)**. Ultimately, this analysis allowed for the determination of whether the *in vitro* consortia accurately captured key aspects of naturally occurring cyanobacterial microbiomes **(Fig. 1B-D)**.

**Assembly independent marker gene analysis of shotgun metagenomic data**

Shotgun metagenomic sequencing was applied to source environmental samples and co-cultures at the 84-day endpoint to recover metagenome-assembled genomes (MAGs) and analyze MAG-encoded functions. Due to the high complexity and low MAG recovery from source samples, we also employed an assembly-independent marker gene analysis using ribosomal protein L6 OTUs to compare diversity between source samples and endpoint co-cultures. Marker gene analysis allowed a more comprehensive assessment of taxonomic diversity, mitigating data loss that occurs during assembly and binning, particularly in highly complex communities. Given the lower recovery of genomic bins from source environmental samples compared to cyanobacterial cultures, and the use of different sequencing approaches, we opted for an assembly-independent marker gene analysis method. This approach allows for a more accurate comparison of diversity and taxonomic composition between environmental samples and cyanobacterial community cultures, drawing on findings from previous studies (Olm 2020) which highlighted the limitations of 16S rRNA genes and the advantages of ribosomal proteins for species discrimination (**Table S8 and S9**). This approach allows the identification of operational taxonomic units (OTUs) directly from shotgun metagenomic read data and circumvents the expected data losses that occur during short read assembly and genomic binning of samples with high species complexity. Briefly, we used SingleM v0.13.2 [43] to search metagenomic reads from each sample against the singleM database v3.0.5 [44] containing 59 conserved single copy marker genes from bacteria and archaea. Search was implemented using the singlem pipe command with default parameters. Identified marker genes were labeled with GTDB taxonomy using the pplacer algorithm for phylogenetic placement. Subsequently OTUs were generated by clustering marker sequences identified across all samples at 95% nucleotide identity using the following command: singlem summarise --input-otu-tables [All singleM pipe outputs] --cluster --cluster-id 0.95 --wide-format-otu-table [output].

Statistical analysis of singleM single copy marker gene OTUs was conducted in R v4.2.3 (See Code Availability). We initially evaluated three potential marker genes with demonstrated potential for species level identification and clustering [45] to conduct the subsequent diversity and taxonomic analysis: ribosomal protein L6 (rpL6), ribosomal protein S3 (rpS3), and ribosomal protein S9 (rpS9). All genes evaluated exhibited a similar level of OTU recovery and predicted similar taxonomic composition across all samples, thus we subsequently chose rpL6 to conduct the full analysis. Samples used in our analysis were subset to a set of 108 cyanobacterial community cultures and 13 source environment samples that were used in MAG resolved metagenomic analysis (See Below). Following sample subsetting OTUs with no counts in any remaining sample were removed resulting in a total final set of 8,156 rpL6 OTUs (**Table S8**). Relative abundance of phylum level taxa was calculated as the fraction of total counts assigned to OTUs grouped at the phylum taxonomic level per sample. Phyla representing less than 1% of the total were collectively categorized under "Other". For clarity, specific phylum names were modified, with "Patescibacteria" being renamed to "CPR". Relative abundance was averaged across all samples displayed in respective plots. Plots were constructed using the ggplot package [34] in R.

Alpha-diversity analysis was conducted on rarefied OTU counts. Briefly, OTU counts across all samples (n = 121 samples) were rarefied using the GUniFrac package in R [35] to the depth of the sample with the lowest number of counts (965 counts). Richness and Shannon diversity metrics were calculated using the microbiome package in R [46] The means of richness and Shannon diversity were compared between all source samples grouped by sampling location using analysis of variance (ANOVA) followed by pairwise comparisons using Tukey's multiple comparisons of means implemented in R with comparisons reporting a FDR ≤ 0.05 being significant. The means of richness and Shannon diversity were compared between all source and all culture samples using Welch's two Sample t-test implemented in R with comparisons reporting a $P \leq 0.05$ being significant. The means of richness and Shannon diversity were compared between all cyanobacterial culture samples grouped by sampled location using analysis of variance (ANOVA) followed by pairwise comparisons using Tukey's multiple comparisons of means implemented in R with comparisons reporting a FDR ≤ 0.05 being significant.

To explicitly assess the dissimilarity of the non-cyanobacterial community composition between environmental source samples and cultures we removed rpL6 OTUs from the dataset that were taxonomically classified in the phylum cyanobacteria and subsequently assessed the beta-diversity of samples using Aitchison distance. Briefly, we first removed rpL6 OTUs classified as cyanobacteria and those with positive counts in < 2 samples (n = 1468 OTUs remaining). We subsequently performed zero imputation on the filtered rpL6 OTU count data using the zCompositions package [41] in R. Centered log-ratio (CLR) transformation was then applied using a custom R function (See Code Availability). Aitchison distance between samples was then calculated as the Euclidean distance between CLR transformed samples using the vegan package [38] in R. The marginal influence of sample type (cyanobacterial culture or source environment) and location (Discovery Bay, Eel River, or Strawberry Creek) were estimated using permutational analysis of variance implemented in the vegan package in R [38] using a model of the form: `Distance_Matrix ~ sample_type + location`. Model terms were assessed marginally using 999 permutations. Principal component analysis was performed in R on centered and scaled CLR transformed rpL6 OTU count data to display community dissimilarities. PCA was chosen for this data display to better preserve the true distances between samples, specifically the large distances between the environmental source environment and cyanobacterial culture samples.

To evaluate the over/under enrichment of bacterial taxonomic orders in cultured cyanobacterial communities relative to source environmental samples we applied a permutation-based method where the observed frequency of taxa in cyanobacterial communities was compared to the frequency of taxa in communities generated through random sampling of all possible taxa identified in both cyanobacterial and source communities. Briefly, we first assessed the number of unique rpL6 OTUs identified in each of the 108 cyanobacterial community samples used in this analysis (**Table S8**). We then created a set of rpL6 OTUs that were non-cyanobacterial and detected in at least one sample across the set of 121 cyanobacterial community and source environment samples (7,773 rpL6 OTUs). We then generated 10,000 permuted data sets of the 108 cyanobacterial community samples where for each sample in a permuted dataset we selected a number OTUs from the total set of 7,773 randomly without replacement equal to the original number of unique rpL6 OTUs identified in that sample (e.g. if a sample had 20 unique OTUs we randomly selected 20 OTUs into its permuted version). This was done such that permuted sample sets had the same OTU richness as the original cyanobacterial community samples. We then aggregated OTUs at the order level and calculated the number of times each order was observed in the set of original 108 samples as well as in each set of the 10,000 permuted samples. Then for each taxonomic order we generated two p-values, one for over enrichment and one for under enrichment, by quantifying the number of times an order appeared more often or less often in the set of permuted samples relative to its observed frequency in the original set of 108 cyanobacterial communities divided by the number of permutations. A combined, two-sided, p-value was generated by taking the minimum of the two p-values and multiplying by 2. A pseudo z-score for each order was generated by subtracting the mean number of times an order was observed across all permuted data sets from the number of times it was observed in the original set of 108 cyanobacterial communities divided by the standard deviation of the number of times an order was observed across all permuted data sets. The p-values for each order were corrected for multiple testing using FDR and comparisons reporting a FDR ≤ 0.05 were considered significant (**Tables S8 and S9**; Also see Code Availability).

**Metagenome assembly, binning, bin de-replication**

Raw reads were processed to remove Illumina adaptors and phiX sequences using BBduk (https://sourceforge.net/projects/bbmap/) and quality-trimmed with Sickle [47]. Each sample was individually assembled using idba_ud( [48], or when idba_ud failed metaspades [48, 49] to generate DNA scaffolds. Scaffolds shorter than 1000 bp were excluded from further analysis. Scaffolds larger than 5000 bp were then subjected to overlap based reassembly using COBRA v50 [50]. Contigs extended and validated by COBRA were non-redundantly re-incorporated into the assembly. Gene prediction was conducted using prodigal v2.6.3 [51] in metagenome mode, and subsequent annotation was performed through a blast search against the uniprot and kegg [47,52, 53] databases. MAG binning was performed on each sample individually by grouping scaffolds longer than 2,500 bp based on sequence compositionality and differential coverage. Cross-mapping of reads from all samples was carried out using bowtie2 [54] to generate differential coverage information for binning. Samples were then independently binned using four algorithms: MetaBat2, maxbin2, CONCOCT, and vamb [55–58]. The best MAGs across the four methods for each sample were selected using DasTool [59]. MAGs were then evaluated for completeness and contamination using checkM [59, 60], which utilizes a single copy gene inventory method. Only MAGs with an estimated completeness of ≥60 % and estimated contamination of ≤5 % were retained. MAGs of sufficient quality across all samples were de-replicated at the species level and representative species MAGs were selected using dRep [61]. MAGs were considered the same species if ANI was ≥ 95 % across ≥ 10 % of the MAG length. Taxonomic placement of MAG was performed using the GTDB-tk [62] classify workflow against the GTDB-R214 taxonomy. Gene prediction was conducted again on non-redundant species-representative MAG using prodigal v2.6.3 [51] in single genome mode. This resulted in a final non-redundant MAG set of 537 MAGs representing all samples in the study. Also see **Table S7.**

**MAG abundance mapping and diversity analysis**

Sequencing reads from each sample were mapped to a concatenated FASTA file of all 537 dereplicated genomic bins using Bowtie2 [54]. Subsequently, CoverM v0.6.1 [63] was used to quantify the number of reads mapped, the coverage, and the breadth of coverage for each MAG. Breadth is defined as the proportion of a genomic bases in a MAG covered by at least one mapped read. Mapped reads were discarded from this analysis if < 50 bp of a read mapped, the read mapping identity was < 97%, or if both reads in a pair did not map in proper orientation and within the estimated sequencing insert length. Finally, for a MAG to be considered as present in a sample we required that the MAG have at least 100 properly mapped reads from that sample and that $\log_2$(Observed Breadth / Expected Breadth) ≥ -1. Expected breadth for a MAG in a sample was calculated as: Expected Breadth = $1 - e^{-0.883 * coverage}$. If a MAG did not pass these filtering criteria its counts and coverage were set to 0 for a sample. These filtering criteria were applied as they significantly reduce the frequency at which MAGs that are not truly in a sample are identified in a sample due read recruitment and mapping to homologous regions shared by multiple MAGs.

Following mapping of sample reads to MAGs we initially identified and filtered out samples from subsequent analysis that did not meet the following criteria: culture samples where ≥ 50% of reads did not map to assembled MAGs (n = 1 sample), culture samples where the expected host cyanobacterium did not comprise ≥ 90% of the cyanobacterial community fraction (n = 11 samples), culture samples that were determined to be mis-labeled following sequencing (n = 1 sample). Of the 134 samples submitted for sequencing 13 source environment samples and 108 cyanobacterial culture samples were utilized for downstream analysis (n = 121 total samples). This set of filtered samples was also used in all other analyses conducted in this study.

Relative abundance of phylum level taxa was calculated as the fraction of total counts assigned to MAGs grouped at the phylum taxonomic level per sample. Phyla representing less than 1% of the total were collectively categorized under “Other”. For clarity, specific phylum names were modified, with “Patescibacteria” being renamed to “CPR”. Relative abundance was averaged across all samples displayed in respective plots. Plots were constructed using the ggplot package [34] in R.

To assess beta-diversity, we calculated Aitchison distances for two datasets: (1) non-cyanobacterial community composition in environmental source samples and cultures (n = 121), and (2) cyanobacterial community cultures alone (n = 108). In both cases, MAGs present in fewer than two samples were removed (501 MAGs remaining for dataset 1; 291 for dataset 2), and zeros were imputed using the zCompositions package [41] in R. CLR transformation was applied, followed by Aitchison distance calculations using vegan [38]. For the environmental vs. culture dataset, we used permutational analysis of variance to test the influence of sample type (culture vs. source environment) and location (Discovery Bay, Eel River, or Strawberry Creek) on community dissimilarity (999 permutations). The marginal influence of sample type and location were estimated using a model of the form: `Distance_Matrix ~ sample_type + location`. Principal component analysis was performed in R on centered and scaled CLR transformed MAG coverage data to display community dissimilarities. PCA was chosen for this data display to better preserve the true distances between samples, specifically the large distances between the environmental source samples and cyanobacterial culture samples. For cyanobacterial culture-only samples, we assessed the effects of host strain, inoculation source location, passage rate, and their interactions using PERMANOVA (9,999 permutations) using a model of the form: `Distance_Matrix ~ host_strain * location * passage_rate`. Model terms and interactions were assessed sequentially using 9,999 permutations. Uniform manifold approximation (UMAP) implemented with the umap package [64] in R was performed on CLR transformed count data to display relationships between samples. UMAP was chosen for this purpose as it provides better recovery of sample clusters and sub-cluster structure, allowing the nested nature of covariate influences to be visible.

**Population level single nucleotide variant analysis**

We evaluated the strain-level similarity of MAGs between cyanobacterial cultures to: (i) Estimate the frequency at which identical strain sharing occurred between cultures, which may indicate cross-contamination; (ii) Evaluate the impact of the inoculum source and cyanobacterial host strain on the microdiversity of individual MAGs across our cyanobacterial communities. To evaluate between-culture strain-level similarity the population level single nucleotide variants (SNVs) were compared for each MAGs between samples using inStrain v1.6.4 [65]. Briefly, we first profiled the SNVs of MAGs in each of the 108 samples using inStrain profile as follows: `inStrain profile [sample_bam_file] [genome_scaffolds] -o [inStrain_output] -p 64 -s [genome_scaffold2bin_file] --min_snp 9999 --skip_mm_profiling`. We applied inStrain defaults using a minimum coverage of 5X to conclusively identify an SNV in a sample. Subsequently we used inStrain compare in database mode to produce an all-vs-all comparison of the MAGs across the 108 SNV profiles generated as follows: `inStrain compare -i [all_108_instrain_profile_outputs] -o [inStrain_compare_output] -p 64 -s [genome_scaffold2bin_file] --database_mode`. We report that this analysis has significantly higher genomic coverage requirements relative to genomic bin recovery and quantitative community compositional analysis. Despite these technical issues we evaluated at least 1 inStrain two-sample comparison for 133 non-host cyanobacterial MAGs in our study (n = 3088 total comparisons). This set included 13 core microbiome species MAGs (n = 555 total comparisons). Subsequent statistical analysis was conducted in R v4.2.3 (**Table S12,** See Code Availability).

To evaluate the frequency at which identical strain sharing occurred between cultures, and potential cross-contamination, we considered a MAG to be identical between samples if its population ANI (popANI) between two samples was ≥ 99.999% based on previously published metrics [65]. We specifically evaluated identical strain sharing occurring between cultures inoculated from distinct environmental sources, as identical strains occurring in cultures inoculated using geographically distinct source material would be probabilistically more indicative of cross-contamination. We found that across a total of 176 two-sample comparisons (n = 11 MAGs; n = 6 core microbiome MAGs) between samples inoculated from different source environments, 0 comparisons were above the popANI threshold to indicate identical strain sharing. Thus, we concluded that although identical species were recovered from geographically distinct source inocula, the species recovered were in fact different strains that were not likely directly transferred between cultures by cross-contamination.

The impact of sharing the same cyanobacterial host strain or source environment on pairwise MAG popANI was modeled using beta regression. Briefly, popANI values for comparisons involving any of the 4 host cyanobacterial species MAGs were removed from the dataset (n = 3,088 comparisons evaluated). As some popANI values were equal to 1 (n = 20 comparisons) a small constant (0.0000001) was subtracted from all popANI values to enable beta regression modeling. popANI was modeled as a function of the inoculation source location being the same or different and the cyanobacterial host strain being the same or different for all 3,088 two sample comparisons using beta regression implemented using the betareg package [66] in R with the following model form: popANI ~ same_location + same_host. Estimated marginal mean popANI values were calculated using the emmeans package in R [40, 65]. Statistical significance for pairwise comparisons between estimated marginal means was evaluated using the multcomp package [67] in R. Tukey's method was applied for comparing multiple means with adjusted $P \leq 0.05$ considered significant. Also see Code Availability.

**Functional Annotation and Core Microbiome Functional Enrichment Analysis**

Predicted proteins in all 537 MAGs were annotated using kofamscan [67, 68] with KOfam HMM set r02_18_2020. Only HMM hits exceeding score thresholds set individually for each HMM were used for downstream analysis, with the highest-scoring annotation selected when multiple significant hits were found. In addition, the pipeline METABOLIC [69] was used to annotate predicted proteins using the function METABOLIC-G.pl and default settings. The CANT-HYD HMM database and hmmsearch (HMMer v. 3.3) with the `--cut_nc` parameter were used to search for genes involved in alkane degradation [70].

To determine metabolic functions enriched in core microbiome MAGs (n = 25) relative to all remaining MAGs (n = 512), Fisher tests were conducted to evaluate the over/under enrichment of each identified KO using the fisher.test function in R v 4.2.1. The p-values were adjusted to account for multiple testing using false discovery rate with the p.adjust function in R, with FDR ≤ 0.05 considered significant. KOs were counted using presence/absence per MAG. Two sets of tests were run on KOs. The first set of tests used all MAGs (n = 537 MAGs) and all KOs that appeared in more than 10 MAGs (n = 4264 KOs). The second set of tests was constrained to MAGs in the phyla Pseudomonadota, Bacteroidota, and Spirochaetota (n = 391 MAGs), and KOs that were found in over 10 MAGs (n = 3749 KOs). As the core microbiome had a highly constrained taxonomic composition, the constrained test was used to evaluate if differential enrichment of KOs was an artifact of taxonomic bias (e.g. was due to functions encoded in taxa not present in the core microbiome) or if the same functions were consistently evaluated as differentially enriched in the core microbiome even when taxonomy was constrained in the analysis. KOs were assigned to broader functional categories as in Diamond et al. [71].

Differential over/under representation analysis of functional categories was performed using the fisher.test function in R v 4.2.1. The p-values were adjusted to account for multiple testing using false discovery rate with the p.adjust function in R, with FDR ≤ 0.05 considered significant (**Tables S16-S18).**

**Identification, curation, and analysis of putative mobile genetic elements**

We identified contigs predicted to be MGEs within the 537 MAGs assembled in this study using geNomad [72] with the parameters: `--end-to-end, --cleanup, --enable-score-calibration, and --splits 8`. Contigs predicted to be plasmids by geNomad were filtered prior to downstream analysis to only include contigs with a FDR ≤ 0.05, and a length over 12 kb. A minimum contig length of 12 kb was chosen as the plasmid prediction accuracy of geNomad has been reported to be ≥ 90.0% for contigs above this length [72]. Predicted plasmids were assigned to MAGs using a guilt-by-association approach where the genomic bin a plasmid contig was found in was considered the putative host. The proportion of contigs predicted to be plasmids per MAG was compared between groups of MAGs assigned to the core, auxiliary, and source microbiome groups using Dunn's test implemented in the dunn.test package in R. The proportion of plasmid contigs per MAG was chosen as the comparison metric to control for differences in total contig number per MAG across all MAGs analyzed. All plasmid contigs were searched against the IMGPR plasmid database using *BLASTn [73]* **(Tables S18-S20)**. HMMs from Conjscan [74] and Pfam [75] were used with hmmsearch (HMMer v. 3.3) to identify genes related to conjugation and mobilization, and replication, respectively, in the plasmid contigs. Definitions for mobilizable or conjugative plasmids were taken from Cury et al. 2020. A plasmid was identified as mobilizable if it contained any of the MOBX genes. A plasmid was identified as conjugative if it contained the relaxase (T4CP) and a ubiquitous ATPase (virB4), as well as three other components of a conjugative system (e.g., virB8, virB9, and virB10). Only HMM hits with an e-value less than 0.001 were considered.

Additional circular element search, curation, and analysis was performed on all core microbiome MAGs (n = 25 MAGs). Briefly, the *de novo* assembly function of Geneious Prime 2023.0.3 (Biomatters Ltd) with default parameters was used to perform overlap assembly on all contigs from each core microbiome MAG. We subsequently evaluated the circularity of assembled contigs in each MAG by searching for overlaps between the first and last 500 bp of each assembled contig larger than 10 kb using BLAST v2.10.1. This effort identified a single putatively circular *de novo* assembled contig (Plasmid_1) in a *Gemmobacter* MAG of the core microbiome (unicom2021_Dbay_Inner_A7120_BW_A_idba_concoct_16). Circularity and *de novo* assembly joins of this contig were verified by mapping of sample reads to the contig using Bowtie2 [54] and visually determining that mapped paired reads spanned all assembly joins and the ends of the contig. Following identification of a circular plasmid in a *Gemmobacter* MAG of the core microbiome we attempted to manually identify additional circular plasmids in the 4 non-representative MAG bins that were recovered during our assembly and binning of samples (See Above), that shared species identity with *Gemmobacter* MAGs of the core microbiome, but were not chosen as the representative MAG bins for those species by dRep. Although the representative MAG bins for a species demonstrate the highest overall quality across a set of metrics, alternative MAG bins recovered for the same species may still have individual contigs with better assembly. By applying the same *de novo* assembly, overlap identification, and read mapping procedure to the 4 additional *Gemmobacter* bins not in our species representative bin set, we identified one additional circular plasmid (Plasmid_2) in a *Gemmobacter* MAG of a different species than Plasmid_1. A large (407 kb) putative plasmid was detected in a Rhizobiales MAG (*StrawCreek_S_L0902_W_A_idba_concoct_4*) using de novo assembly and sequence overlap identification. Although this element was not circularized, it encoded plasmid-specific replication factors (*repABC*), a complete conjugal transfer system (*TraI*, *trb*), and gene clusters for succinoglycan and alginate biosynthesis **(Table S23, Fig. 6D)**. Genes on these three plasmids were annotated with prodigal v2.6.3 [51], and functional annotations were predicted using kofamscan [68] with KOfam HMM set r02_18_2020 (**Table S23**).

Assessment of functions putatively enriched on MGEs associated with MAGs of the core microbiome was conducted by repeating KEGG KO enrichment analysis on all 537 MAGs assembled in this study after removing contigs identified as plasmids by genomad. Briefly, we first removed all 850 contigs classified as predicted plasmids by genomad from MAGs. We then re-quantified the presence/absence of KEGG KOs in each MAG with predicted plasmid contigs removed. All KOs used in the original full enrichment analysis (See Above; n = 537 MAGs) were re-evaluated for statistical over/under enrichment in core microbiome MAGs relative to non-core microbiome MAGs using the *fisher.test* function in R v 4.2.1. The p-values were adjusted to account for multiple testing using false discovery rate with the *p.adjust* function in R, with FDR ≤ 0.05 considered significant. We then identified the set of KOs that were significantly enriched in the original core microbiome analysis, but were no longer significantly enriched in the core microbiome analysis with MGE contigs removed. A relative change between the two analyses for each KO was calculated as: Fold Change in Odds Ratio = KO_Odds_Ratio_Original / KO_Odds_Ratio_MGE_Removed.

**Identification and evaluation of unique KEGG functions in host cyanobacterial species**

Using the KEGG KO annotations of all genomes in this study (See Above) we identified 1886 KOs that could be found in one or more of the 4 cyanobacterial host species genomes evaluated (**Table S24**). From this set we identified 447 KOs that were unique to only one host cyanobacterial species genome. Summary statistics on unique KOs by functional category were calculated in R v 4.2.1 and are available in **Table S25**. The over- or under-enrichment of unique KOs across all genomes within each functional category was evaluated using Fisher's exact test using the fisher.test function in R v 4.2.1. The Fisher test was conducted for each functional category independently using a 2 X 2 contingency table as follows:

| | |
|---|---|
| Unique KOs in Functional Category | Unique KOs Not in Functional Category |
| Non-Unique KOs in Functional Category | Non-Unique KOs Not in Functional Category |

The natural log transformed odds ratio produced by the Fisher test function was reported. The p-values across all Fisher tests were adjusted to account for multiple testing using false discovery rate with the p.adjust function in R, with FDR ≤ 0.05 considered significant (**Table S25**). Plots were constructed using the ggplot package [34] in R.

### Supplementary Results

#### Inoculation of model cyanobacterial strains with bacteria from diverse freshwater sources established stable *in vitro* communities.

Out of the 180 initial *in vitro* consortia, 108 consortia were retained through the 12-week passaging period, with each consortium retaining its cyanobacterial host species as the dominant organism (≥ 90%). A detailed logistic regression analysis showed that several factors significantly influenced the retention of cultures. Specifically, the species of cyanobacterial host strain was a strong predictor of retention ($P$ = 3.412e-9). The inoculum source location was also a significant factor ($P$ = 1.236e-4), as was the interaction between the host strain and passage rate ($P$ = 0.0122). However, passage rate alone did not have a significant impact on retention ($P$ = 0.203) **(Fig. S2 and Table S2)**.

Among the host strains, *Synechocystis* spp., PCC 6803 (6803), *Leptolyngbya* spp., BL0902 (L0902), and *Nostoc* spp., PCC 7120 (A7120) demonstrated the highest retention rates, whereas *Synechococcus elongatus* UTEX 3055 (3055) and *Synechococcus elongatus* PCC 7942 (7942) showed lower retention rates **(Table S2)**. Consortia inoculated with microbial fractions sourced from the Eel River had the highest retention rates, outperforming those derived from Discovery Bay or Strawberry Creek inocula **(Fig. S2)**. These results suggest that the specific cyanobacterial host and the inoculum source environment strongly influenced the successful establishment of stable consortia.

Among the 72 cultures that were not retained through the full passaging period, qualitative light microscopy analysis revealed the presence of protozoans in a number of cultures. This observation suggests that protozoan predation may have contributed to the failure of these consortia, as predation is a well-documented challenge in dense cyanobacterial cultures exposed to natural microbial communities [27]. Protozoan contamination can rapidly destabilize cyanobacterial cultures, leading to collapse.

For the 108 consortia that were retained, 16S rRNA gene sequencing data were collected at multiple time points throughout the 12-week passaging period (n = 972 samples) to monitor changes in community diversity, taxonomic composition, and stability. Significant changes in both alpha diversity (Shannon index diversity and richness) and beta diversity were observed within the first 14 days of cultivation, indicating substantial shifts in community structure during the early stages of consortia formation **(Fig. 1B-D, Table S3-S6)**. However, after 28 days, the consortia stabilized, and no significant changes were observed in either diversity metrics or taxonomic composition beyond this point **(Table S3-S6).**

To assess the robustness of the consortia, samples were subjected to cryopreservation followed by revival and further passaging. The results showed that cryopreservation and subsequent revival did not significantly affect community composition or diversity metrics **(Table S3-S6, Fig. S3).** No major shifts in community composition were detected, confirming that the consortia were stable and resilient to the freeze-thaw process. These findings demonstrate that the consortia can be cryopreserved without compromising their structure, making them suitable for long-term storage and potential future use.

The final consortia that developed after 12 weeks of cultivation were highly similar to natural cyanobacterial communities in terms of both taxonomic composition and relative abundance **(Fig. 1, Table S6).** Despite the different sources of microbial inocula and the use of various host cyanobacterial strains, the consortia converged on a consistent taxonomic structure, dominated by the same microbial groups commonly observed in natural environments [24, 25]. This suggests that the *in vitro* model system accurately captures key aspects of natural cyanobacterial microbiomes, and that the dominant taxa in these systems are robustly selected during the early stages of community formation, regardless of the inoculum source or host strain.

**Genomic resolution enabled comprehensive characterization of *in vitro* communities**

We performed shotgun sequencing on both the stable *in vitro* consortia at the 84-day time point and their corresponding environmental source samples **(Table S2).** This analysis yielded a total of 537 non-redundant species-level draft MAGs. Based on MIMAG standards, 321 MAGs (59.8%) were classified as high-quality, with a completeness of ≥90% and contamination ≤5%, 216 MAGs (40.2%) were classified as medium-quality, with a completeness of ≥60% and contamination ≤10% **(Table S9)**. These recovered MAGs spanned 17 distinct phyla, with *Bacteroidota* (n = 149 MAGs) and *Pseudomonadota* (n = 133 *Alphaproteobacteria* and n = 106 *Gammaproteobacteria*) being the most prominently represented.

Of the 537 MAGs, 324 species were found in the *in vitro* consortia, and 209 species were exclusive to the environmental source samples **(Fig. 2A).** 84.2% of these MAGs had no species representative in existing genomic databases, with 59 MAGs unclassified beyond the family and 8 MAGs beyond the orderlevel **(Table S9)**. Read-mapping analyses showed that the recovered MAGs provided a strong representation of the *in vitro* communities, with an average of 86.2% of reads mapping back to these MAGs (IQR = 83.3 - 90.3%) across all samples **(Table S2).** By contrast, only 9.3% (IQR = 4.5 - 12.8%) of reads from the source environment samples mapped to MAGs recovered from the source.

**Specific taxonomic groups become enriched in stable *in vitro* communities**

Even though our approach is MAG-centric, many microorganisms in the source environment samples were too low in abundance to be represented by MAGs. Therefore, we applied assembly-independent marker gene analysis to the shotgun metagenomic data to directly compare the composition of the source samples with the stable (84 d) *in vitro* consortia **(Table S11,** See Methods). Alpha-diversity, as measured by richness and Shannon diversity, was significantly lower in the *in vitro* consortia compared to the environmental source samples, indicating strong selective pressures during community formation **(Fig. S4).**

Using permutation-based analysis, we identified significant patterns of taxonomic enrichment. Out of 428 detected order-level taxa, 22 orders were significantly overrepresented, and 59 were underrepresented in the *in vitro* consortia compared to the source environments **(Fig. 2B, Table S8).** Orders within *Pseudomonadota* and *Bacteroidota* were the most consistently overrepresented, and orders within Candidate Phyla Radiation (CPR) and *Omnitrophota* were underrepresented. These results suggest that, regardless of the source inocula or host strain differences, the *in vitro* environment selectively enriched specific taxa, and many taxa from species-rich environments such as Strawberry Creek, particularly CPR organisms, were not sustained in the final communities **(Fig. S2, S4).**

The statistical significance of over- and underrepresented taxa in *in vitro* consortia compared to the source samples was determined using permutation-based analysis. A total of 10,000 permutations were performed to determine the Z-scores for each taxonomic order **(Table S8).** Orders with a false discovery rate (FDR) of ≤ 0.05 were considered statistically significant. Orders such as *Rhizobiales*, *Sphingomonadales*, and *Rhodobacterales* showed significant enrichment, CPR and *Omnitrophota* showed significant underrepresentation.

**Identical species were selected into communities from geographically distinct sites**

To assess whether the presence of identical core and auxiliary species across our communities resulted due to selection from diverse environmental sources or alternatively from cross-contamination, we analyzed population level single nucleotide variants (SNVs) of species populations between samples. We employed inStrain (See Methods) to compare these SNVs and conducted 3088 two-sample comparisons for 133 species, which included comparisons of 13 core microbiome species (**Table S13-S14**). Between *in vitro* communities with different source inocula, no pairwise average nucleotide identity (popANI) comparisons showed identical strain sharing (n = 0/176 comparisons). Thus, even though the same species could be detected in communities derived from different source inocula, the strain populations of these species between samples had sufficient nucleotide divergence to indicate that they were derived from different original populations (e.g. from their respective source environments). Furthermore, we observed that the popANI between the same species from different communities was significantly influenced by the identity of the cyanobacterial host (**Fig. S8, Table S13-S14**). We found that divergence in popANI was significantly larger when a species was compared between communities with different source inocula (Estimate = 1.172, $P < 2.00e^{-16}$; beta regression) as well as with different cyanobacterial hosts (Estimate = 0.097, $P = 0.008$; beta regression). These results lend strong support to the hypothesis that identical species, from distinct strain populations, were independently selected into our *in vitro* communities from geographically disparate water sources, rather than being introduced through cross-contamination. Additionally, the identity of a community's cyanobacterial host plays a significant, albeit secondary, role in shaping these populations. This selection pressure results in strain populations associated with the same host species exhibiting greater genetic similarity.

**Core microbiome MAGs are enriched in putative plasmid elements**

Extrachromosomal replicons and plasmids have been shown to encode functions that mediate phototroph-heterotroph symbioses, specifically plant-microbe interactions [76]. Thus, we assessed if there were differences in the frequency and distribution of putative plasmid elements associated with the MAGs resolved in our study. We identified 850 contigs across all species-representative MAG bins (n = 537 MAGs) as putative plasmids (**Table S21**). Most were linear fragments (98%), and 17 were complete circular elements. We additionally manually curated two additional large (>130 kb) circular elements each linked to a *Gemmobacter* MAG within the core microbiome resulting in a total of 3 confirmed circular elements in core microbiome MAGs (**Table S23**). Subsequently we quantified the presence of plasmid specific functions on all putative plasmid contigs and found that 15% encoded proteins with plasmid-specific replication domains and 26% encoded proteins for plasmid mobilization (**Fig. S10A**), typical frequencies for fragmented environmental plasmid sequences [77]. Although environmentally derived plasmid sequences remain under sampled, alignment of putative-plasmids recovered in this study to the IMG/PR database [73] found 81 plasmid contigs that matched (≥ 95% Identity and ≥ 20% coverage) IMG/PR plasmids or plasmid fragments, including 13 associated with core microbiome MAGs **(Table S21)**. We also report that the majority (80%) of matches to IMG/PR were to plasmids recovered from metagenomic datasets, particularly from aquatic environments.

We found that core microbiome MAGs had a significantly higher proportion of contigs predicted to be putative plasmids than MAGs of auxiliary species (Dunn Test = 3.7, $P = 0.0003$) and MAGs detected only in source environment samples (Dunn Test = 6.5, $P = 0$) (**Fig 6A**). As bacteria from the order *Rhizobiales* are known to encode large numbers of extrachromosomal plasmids [78, 79] we also evaluated the enrichment of putative plasmids between MAG groups using only *Rhizobiales* MAGs and found that plasmid enrichment patterns were preserved with core *Rhizobiales* MAGs containing a significantly larger proportion of contigs predicted to be putative plasmids than auxiliary (Dunn Test = 3.0, $P$ = 0.0036) or source (Dunn Test = 2.4, $P$ = 0.0253) environment MAGs **(Fig. S10D)**.

#### Comparative analysis of unique functions between cyanobacterial host species

Differences in metabolic capabilities among the cyanobacterial host species used in this study likely influence microbiome recruitment and potential cross-feeding dynamics. While our primary focus was to characterize these communities under standard laboratory conditions optimized for axenic cyanobacterial growth, we recognized the value of systematically evaluating host-specific functional differences, despite experimental dissection of these differences being beyond the scope of the current study.

Across all host genomes, we identified 1,886 KEGG ortholog (KO) functions, of which 447 were unique to a single host species (Table S24). The number of unique KEGG functions per genome (**Fig. S11A, Table S24**) was significantly correlated with the number of protein-coding genes (r = 0.98; P = 0.016, Pearson correlation). These unique functions spanned 17 broad functional categories. The largest, labeled “Other” (n = 231 genes), encompassed diverse auxiliary metabolic functions not clearly assigned to a specific category (**Fig. S11B**), suggesting that genome expansion in these species is associated with acquisition of accessory metabolic capabilities.

We next evaluated whether any of the 17 functional categories were significantly over- or under-represented among host-unique functions (**Fig. S11C, Table S25**). Core metabolic categories, including Photosynthesis ($_{Log}$OR = -1.06, FDR = .007; Fisher test), Energy Metabolism ($_{Log}$OR = -1.70, FDR = 7.6e-4; Fisher test), and Translation ($_{Log}$OR = -3.35, FDR = 1.4e-9; Fisher test), were significantly under-enriched, consistent with their conservation across cyanobacteria and their essential roles in photoautotrophic growth. In contrast, enrichment of host-unique functions was observed in categories such as Nitrogen ($_{Log}$OR = 1.37, FDR = 1.7e-4; Fisher test) and Phosphonate Metabolism ($_{Log}$OR = 3.83, FDR = 3.0e-7; Fisher test), as well as the broad “Other” category ($_{Log}$OR = 0.80, FDR = 2.1e-14; Fisher test).

Finally, we examined the species distribution and functional identities of host-unique KOs within the over-enriched categories of Nitrogen Metabolism, Phosphonate Metabolism, and the broad “Other” category (**Fig. S11D**, **Table S24**). Within the “Other” category, the proportion of host-unique KOs contributed by each strain was strongly correlated with the number of protein-coding genes in each genome (r = 0.99; $P$ = 0.010, Pearson correlation).

In contrast, unique functions related to Nitrogen and Phosphonate Metabolism were predominantly found in *Anabaena* spp., PCC 7120. These included key KOs involved in nitrogen fixation and the utilization of methylphosphonate as a phosphorus source. These results highlight species-specific metabolic adaptations that may play important roles in shaping microbiome structure and resource exchange under nutrient-variable conditions. They also lay the groundwork for future studies aimed at experimentally testing how variation in nitrogen and phosphorus availability influences microbiome assembly dynamics.

13. Blanc-Garin V, Chenebault C, Diaz-Santos E, Vincent M, Sassi J-F, Cassier-Chauvat C, et al. Exploring the potential of the model cyanobacterium *Synechocystis* PCC 6803 for the

photosynthetic production of various high-value terpenes. *Biotechnol Biofuels Bioprod* 2022; **15**: 110.

14. Rippka R, Stanier RY, Deruelles J, Herdman M, Waterbury JB. Generic assignments, strain histories and properties of pure cultures of Cyanobacteria. *Microbiology* 1979; **111**: 1–61.

15. Lázaro S, Fernández-Piñas F, Fernández-Valiente E, Blanco-Rivero A, Leganés F. pbpB, a gene coding for a putative penicillin-binding protein, is required for aerobic nitrogen fixation in the cyanobacterium *Anabaena* sp. strain PCC7120. *J Bacteriol* 2001; **183**: 628–636.

16. Zeng X, Zhang C-C. The making of a heterocyst in cyanobacteria. *Annu Rev Microbiol* 2022; **76**: 597–618.

17. Koksharova OA, Wolk CP. Genetic tools for cyanobacteria. *Appl Microbiol Biotechnol* 2002; **58**: 123–137.

18. Banerjee M, Raghavan PS, Ballal A, Rajaram H, Apte SK. Oxidative stress management in the filamentous, heterocystous, diazotrophic cyanobacterium, *Anabaena* PCC7120. *Photosynth Res* 2013.

31. Edgar RC. UNOISE2: improved error-correction for Illumina 16S and ITS amplicon sequencing. *bioRxiv* 2016; 081257

32. Yilmaz P, Parfrey LW, Yarza P, Gerken J, Pruesse E, Quast C, et al. The SILVA and 'All-species Living Tree Project (LTP)' taxonomic frameworks. *Nucleic Acids Res* 2013; **42**: D643–D648.

33. Bolyen E, Rideout JR, Dillon MR, Bokulich NA, Abnet CC, Al-Ghalith GA, et al. Reproducible, interactive, scalable and extensible microbiome data science using QIIME 2. *Nat Biotechnol* 2019; **37**: 852–857.

34. ggplot2. https://ggplot2.tidyverse.org/. Accessed 29 Nov 2023.

35. GUniFrac: Generalized UniFrac Distances, Distance-Based Multivariate Methods and Feature-Based Univariate Methods for Microbiome Data Analysis. *Comprehensive R Archive Network (CRAN)*. https://CRAN.R-project.org/package=GUniFrac. Accessed 30 Nov 2023.

36. Website. http://microbiome.github.io.

37. Bates D, Mächler M, Bolker B, Walker S. Fitting linear mixed-effects models Using lme4. *J Stat Softw* 2015; **67**.

38. Community Ecology Package [R package vegan version 2.6-4]. 2022.

39. MuMIn: Multi-Model Inference. *Comprehensive R Archive Network (CRAN)*. https://CRAN.R-

project.org/package=MuMIn. Accessed 30 Nov 2023.

40. Lenth RV. Estimated Marginal Means, aka Least-Squares Means [R package emmeans version 1.8.9]. 2023.

41. Palarea-Albaladejo J, Martín-Fernández JA. zCompositions — R package for multivariate imputation of left-censored data under a compositional approach. *Chemometrics Intellig Lab Syst* 2015; **143**: 85–96.

42. Mallick H, Rahnavard A, McIver LJ, Ma S, Zhang Y, Nguyen LH, et al. Multivariable association discovery in population-scale meta-omics studies. *PLoS Comput Biol* 2021; **17**: e1009442.

43. GitHub - wwood/singlem: Novelty-inclusive microbial community profiling of shotgun metagenomes. *GitHub*. https://github.com/wwood/singlem. Accessed 28 Feb 2025.

44. Woodcroft BJ. Default SingleM reference ‘metapackage’ data.

64. UMAP. <https://CRAN.R-project.org/package=umap.
