## Supplementary Figures for "Model cyanobacterial consortia reveal a consistent core microbiome independent of inoculation source or cyanobacterial host species"

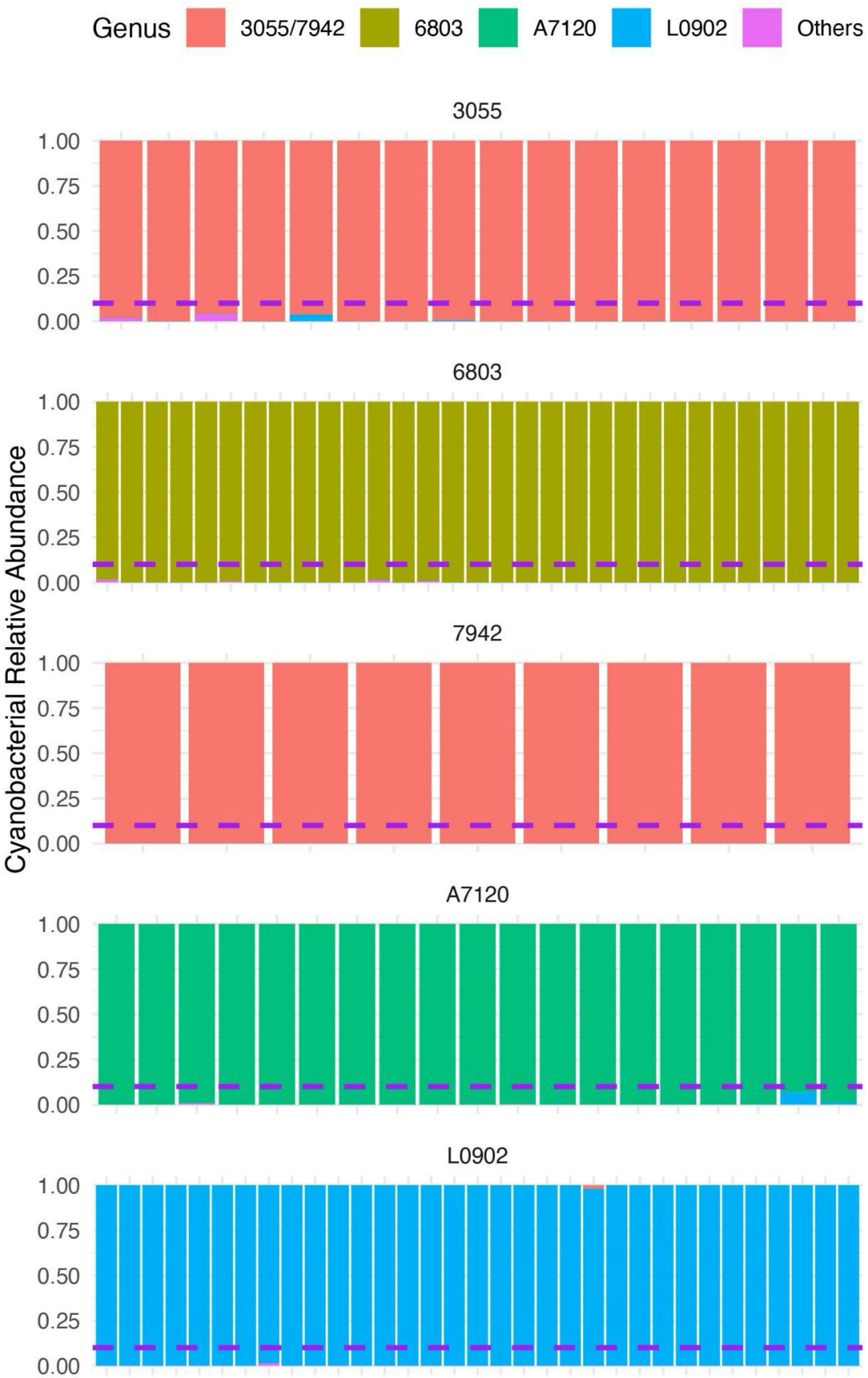


**Supplementary Fig. 1 | Relative abundance of cyanobacterial metagenome-assembled genomes**

**(MAGs) across retained cultures at the 84-day time point**. Genus-level, MAG - based on displaying the relative abundance of cyanobacterial species in co-cultures with specific cyanobacterial host strains (3055, 6803, 7942, A7120, and L0902). Each panel represente communities around different host strain. Bars represent the proportion of each cyanobacterial species within individual samples. The dashed purple line marks the 10% relative abundance.

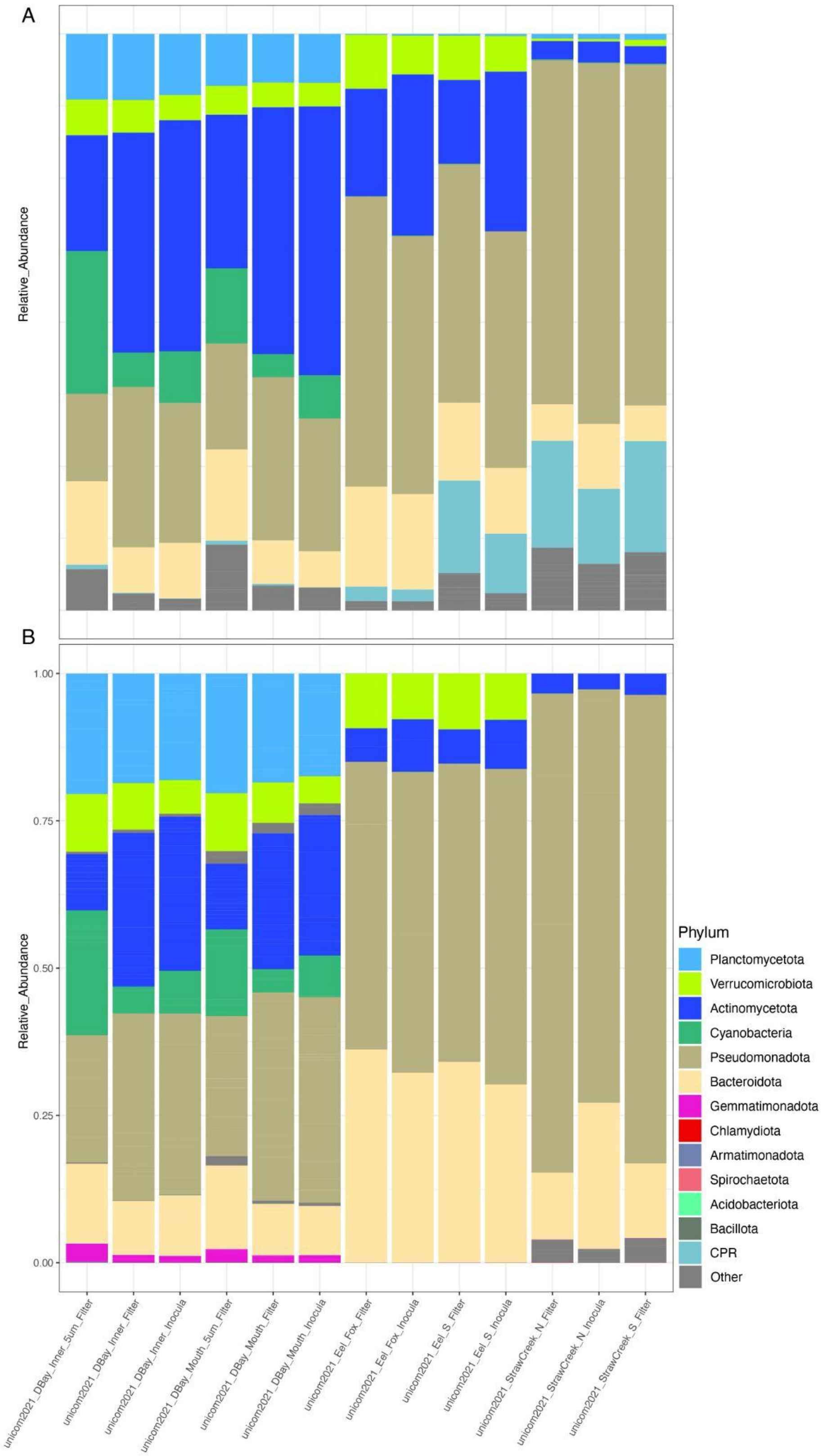


**Supplementary Fig. 2 | Microbial community composition of environmental source microbiomes used for co-culturing with axenic cyanobacterial strains. (A)** Read-based analysis of microbial composition using mapping of reads to 10,166 predicted ORFs of the ribosomal gene marker (rpl6) across six reading frames. **(B)** Phylum-level, MAG-based analysis of microbial communities using mapping of sample reads to 537 species-level MAGs. Taxonomy assignments were conducted using GTDB. A total of 13 samples were examined. The x-axis represents the location and sample type data originated from. “Filter” refers to the combined bacterial biomass on 0.1 and 0.2 μm filters, “Inocula” refers to the biomass dissociated from these filters for culturing, and “5 μm filter” denotes the biomass from the 5 μm filters.

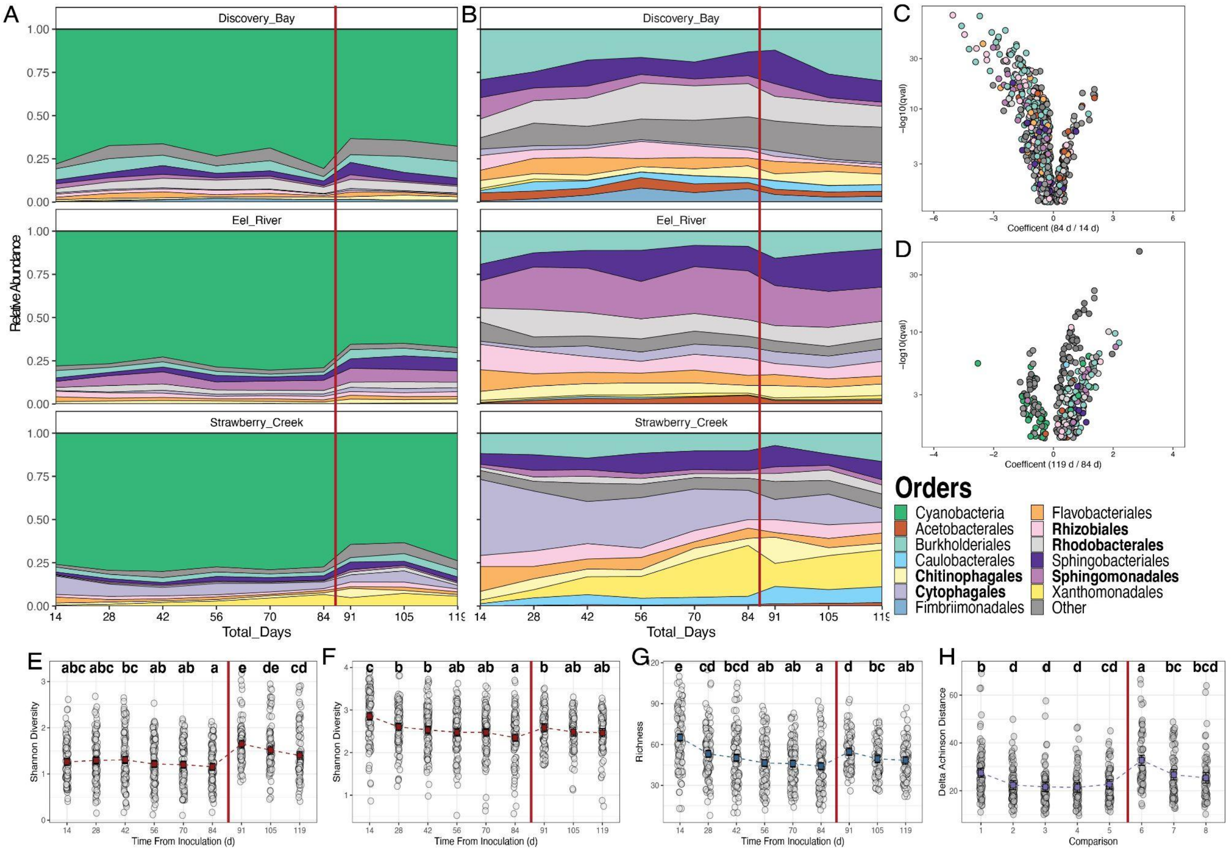


**Supplementary Fig. 3. Stability assessment of *in vitro* communities over the 119-day passaging experiment. (A)** Order-level community composition based on 16S rRNA amplicon sequences, identifying 2126 ASVs over the 119 d time course. The data are presented by location, with a vertical red line marking the division between pre- and post-cryopreservation passaging periods. **(B)** Order-level community composition based on the same dataset but excluding cyanobacterial ASVs to highlight the dynamics of non-cyanobacterial ASVs. **(C, D)** Differential abundance analysis conducted using the MaAsLin2 function in R, applied a linear model to evaluate the relative abundance of 1,557 ASVs (only ASVs with at least 10 positive values across all samples), focusing on selective pressures impacting specific taxa. No minimum thresholds for abundance, prevalence, or variance were set. The significance threshold was established at a p-value of ≤ 0.05, with adjustments for multiple comparisons using the Benjamini-Hochberg method. Fixed effects included General Site, Strain, Passage Rate, and Total Days, with Discovery Bay as the reference level for General Site. Tube Number was incorporated as a random effect. There was strong selection for and against specific order level taxa across all communities as Rhodobacterales and Burkholderiales respectively. **(C)** Day 14 to Day 84: Of the 603 ASVs showing changes in relative abundance, 487 decreased and 116 increased. **(D)** Day 84 to Day 119 (Post-Cryopreservation): Changes were observed in 315 ASVs; 73 decreased and 242 increased in relative abundance. Notably, almost all taxa present before cryopreservation were retained, with no significant decrease in ASV count. **(E)** Observed Shannon diversity of all 2126 ASVs of *in vitro* communities at 14 d intervals over 119 d. **(F)** Observed Shannon diversity, excluding cyanobacterial ASVs, of *in vitro* communities at 14 d intervals over 119 d. **(G)** Observed richness, excluding cyanobacterial ASVs, of *in vitro* communities at 14 d intervals over 119 d. **(H)** Change in Aitchinson distance between previous passage time points for *in vitro* communities at 14 d intervals over 119 d. In panels E, F and G gray dots represent values for each community at respective time points, while in panel H, gray dots depict the change for each community compared to its previous sample. For all metrics

**(E - H)**, colored squares (blue for Richness, red for Shannon Diversity, and purple for Aitchison Distance) indicate estimated marginal means, with error bars representing the 95% confidence intervals (CI) of these means. Statistical significance between time points and comparisons was determined using linear mixed-effects models, with different letters denoting significant differences (False Discovery Rate, FDR ≤ 0.05). A vertical red line marks the division in all plots between pre- and post-cryopreservation passaging periods.

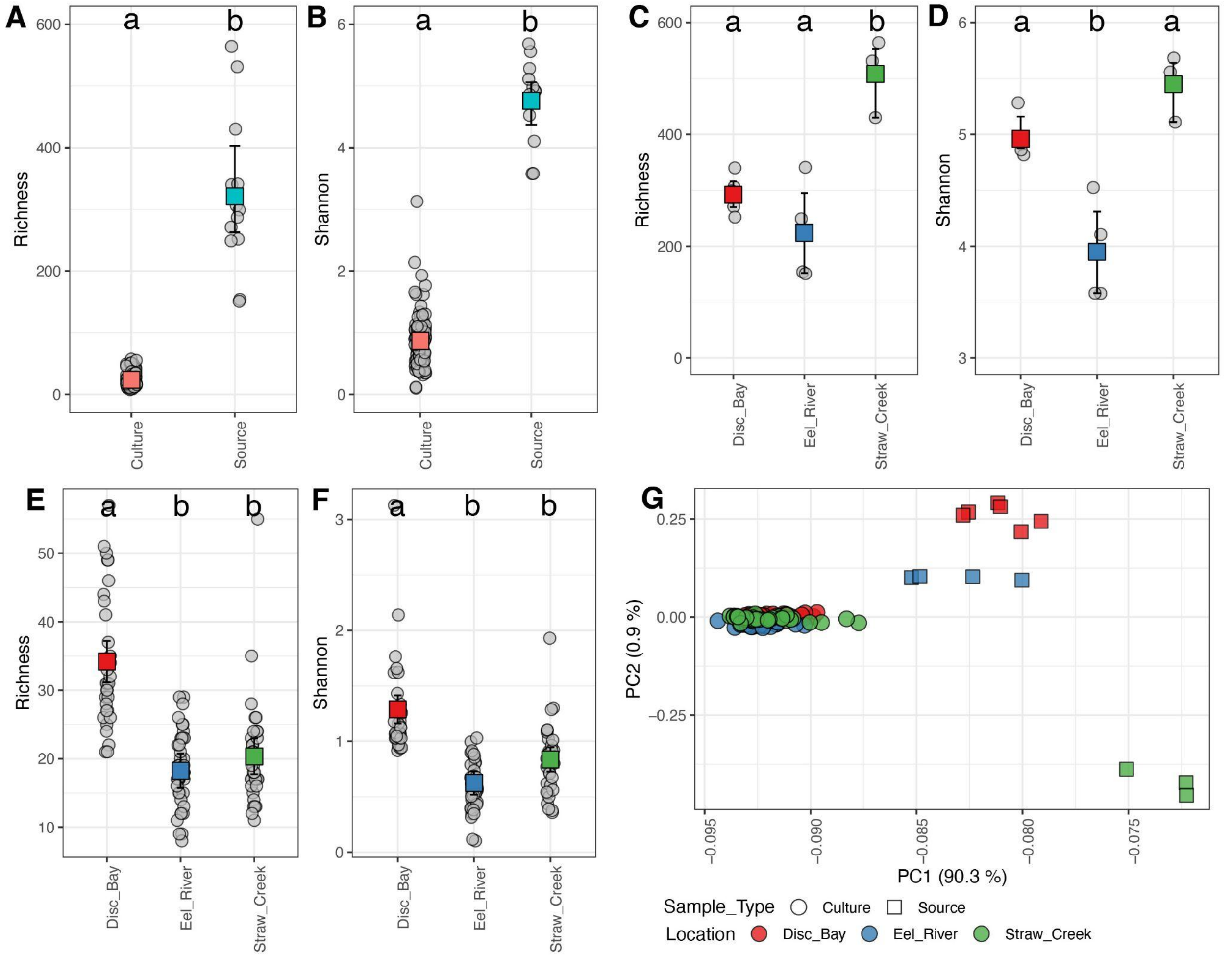


**Supplementary Fig. 4. Variations in microbial community richness, Shannon diversity, and compositional ordination based on rpL6 OTUs. (A, B)** Microbial diversity of all samples by sample type: *in vitro* communities (Culture) had significantly lower richness (mean = 23.76) and Shannon diversity (mean = 0.87) relative to Source samples (richness mean = 321.15, Shannon mean = 4.76). Statistical tests confirmed strong differences (Richness: $t = -8.59$, $df = 12.02$, $p < 0.0001$; Shannon: $t = -20.68$, $df = 13.22$, $p < 0.0001$). **(C, D)** Microbial diversity by source location: Richness and Shannon diversity significantly differed among Source samples from Strawberry Creek, Eel River, and Discovery Bay (Richness: $F = 19.02$, $p < 0.001$; Shannon: $F = 22.77$, $p < 0.001$). **(C)** Post-hoc Tukey HSD tests for Richness showed significant differences between Strawberry Creek and Eel River (mean difference = 284.58, $p < 0.001$) and between Strawberry Creek and Discovery Bay (mean difference = 215.83, $p = 0.002$), while Eel River and Discovery Bay were not significantly different (mean difference = -68.75, $p = 0.25$). **(D)** For Shannon diversity, significant differences were found between Eel River and Discovery Bay (mean difference = -1.02, $p = 0.0012$) and between Strawberry Creek and Eel River (mean difference = 1.50, $p < 0.001$), but not between Strawberry Creek and Discovery Bay (mean difference = 0.49, $p = 0.11$). **(E, F)** Microbial diversity in culture samples by source inoculum: Richness and Shannon diversity varied in culture samples based on their source inoculum from Strawberry Creek, Eel River, or Discovery Bay (Richness: $F = 19.02$, $p < 0.001$; Shannon: $F = 22.77$, $p < 0.001$).**(E)** Post-hoc Tukey HSD tests for Richness showed significant differences between cultures inoculated from Strawberry Creek vs. Eel River (mean difference = 284.58, $p < 0.001$) and Strawberry Creek vs. Discovery Bay (mean difference = 215.83, $p = 0.002$), but no significant difference between Eel River and Discovery Bay (mean difference = -68.75, $p = 0.25$). **(F)** For Shannon diversity, significant differences were observed between Eel River and Discovery Bay (mean difference = -1.02, $p = 0.0012$) and between Strawberry Creek and Eel River (mean difference = 1.50, $p < 0.001$), but not between Strawberry Creek and Discovery Bay (mean difference = 0.49, $p = 0.11$). Squares indicate sample means, error bars indicate 95% confidence intervals, and different letters denote statistically distinct groups. **(G)** Ordination of microbial community compositional dissimilarity. Principal component analysis (PCA) of CLR-transformed rpL6 OTU abundance in all samples (n = 1468 OTUs analyzed, excluding cyanobacteria) reveals strong compositional separation between *in vitro* and source samples. PCA was used to preserve large distances between these groups. Environmental samples (squares) and *in vitro* communities (circles) are clearly distinguished, with symbol colors representing either the bacterial inoculum's geographic origin.

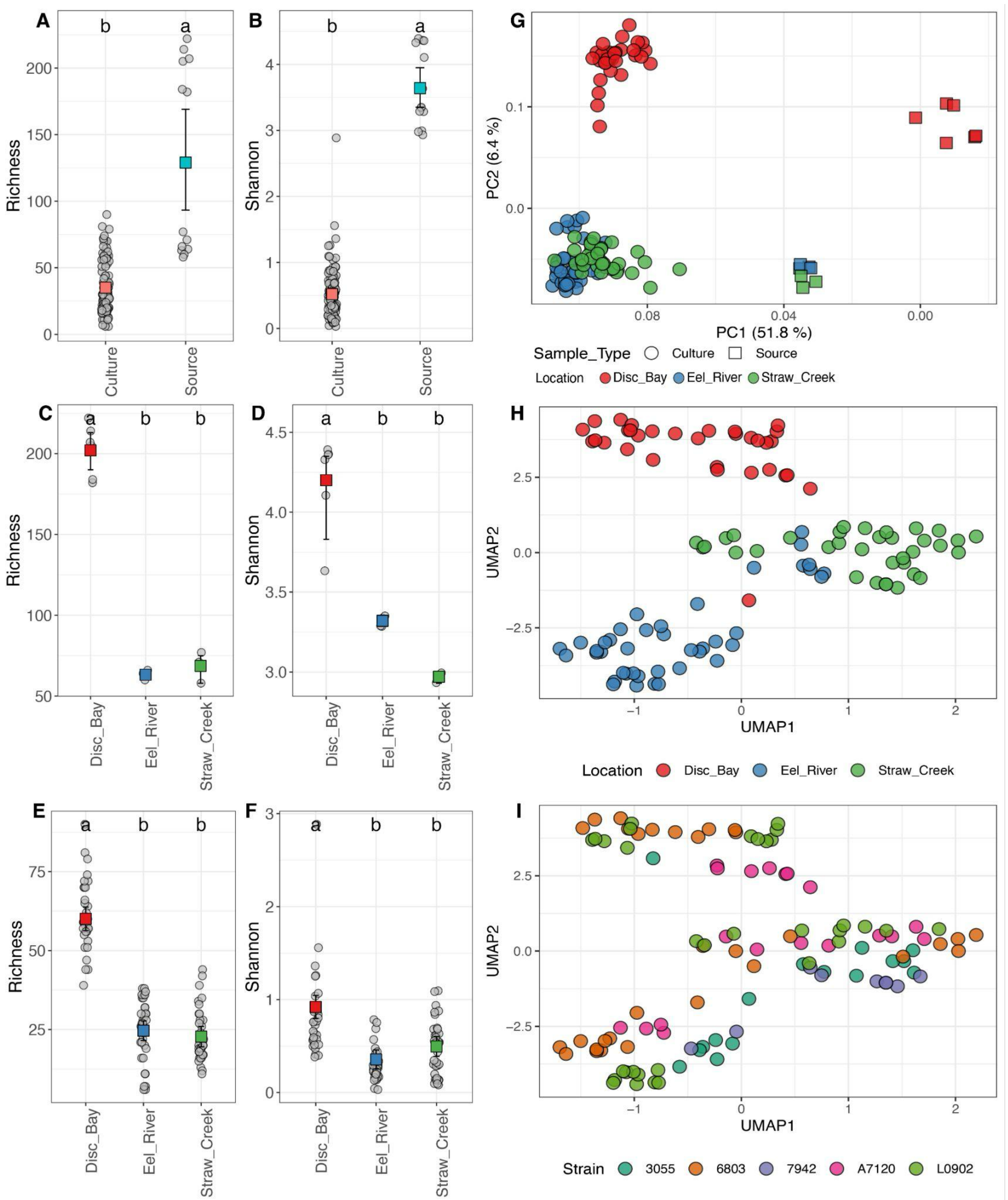


**Supplementary Fig. 5 | Variations in microbial community richness, Shannon diversity, and compositional ordination based on MAGs.**

**(A, B)** Microbial diversity of all samples grouped by sample type: *in vitro* communities (Culture) had significantly lower richness (mean = 35.09) and Shannon diversity (mean = 0.52) relative to Source samples (richness mean = 128.69, Shannon mean = 3.64). Statistical tests confirmed strong differences (Richness: $t = -4.67$, $df = 12.21$, $p = 0.0005$; Shannon: $t = -18.82$, $df = 13.32$, $p < 0.0001$).

**(C, D)** Microbial diversity of source samples grouped by source location: Richness and Shannon diversity significantly differed among Source samples from Strawberry Creek, Eel River, and Discovery Bay (Richness: $F = 200.4$, $p < 0.0001$; Shannon: $F = 41.19$, $p < 0.0001$). **(C)** Post-hoc Tukey HSD tests for Richness showed significant differences between Discovery Bay and both Eel River (mean difference = 139.08, $p < 0.0001$) and Strawberry Creek (mean difference = 133.67, $p < 0.0001$), while no significant difference was observed between Eel River and Strawberry Creek (mean difference = 5.42, $p = 0.83$). **(D)** For Shannon diversity, significant differences were found between Discovery Bay and both Eel River (mean difference = 0.88, $p = 0.0002$) and Strawberry Creek (mean difference = 1.23, $p < 0.0001$), but not between Eel River and Strawberry Creek (mean difference = 0.35, $p = 0.13$).

**(E, F)** Microbial diversity of culture samples grouped by source inoculum: Richness and Shannon diversity varied in culture samples based on their source inoculum from Strawberry Creek, Eel River, or Discovery Bay (Richness: $F = 168.1$, $p < 0.0001$; Shannon: $F = 24.98$, $p < 0.0001$). **(E)** Post-hoc Tukey HSD tests for Richness showed significant differences between cultures inoculated from Discovery Bay vs. Eel River (mean difference = 36.21, $p < 0.0001$) and Discovery Bay vs. Strawberry Creek (mean difference = 38.10, $p < 0.0001$), but no significant difference between Eel River and Strawberry Creek (mean difference = -1.89, $p = 0.67$). **(F)** For Shannon diversity, significant differences were observed between Discovery Bay and both Eel River (mean difference = 0.53, $p < 0.0001$) and Strawberry Creek (mean difference = 0.37, $p = 0.00002$), but not between Eel River and Strawberry Creek (mean difference = 0.16, $p = 0.08$). Squares indicate sample means, error bars indicate 95% confidence intervals, and different letters denote statistically distinct groups. **(G)** Analysis (PCA) of CLR transformed MAGs abundance in all samples. Host cyanobacterial MAGs were excluded from the analysis (n = 501 MAGs analyzed). **(H)** Uniform Manifold Approximation and Projection (UMAP) of CLR transformed metagenomic species abundance in *in vitro* community cultures only. Samples are colored by the source location used to inoculate the community. **(I)** Uniform Manifold Approximation and Projection (UMAP) of CLR transformed metagenomic species abundance in *in vitro* community cultures only. Samples are colored by the cyanobacterial host used in the community. PCA was used to preserve large distances between source and culture samples. UMAP was used to preserve clustering and sub-clustering related to sample covariates in *in vitro* community samples. Environmental samples (squares) and *in vitro* communities (circles) are clearly distinguished, with symbol colors representing the bacterial inoculum's geographic origin or the communities around specific host cyanobacteria. Also see **Table S11**.

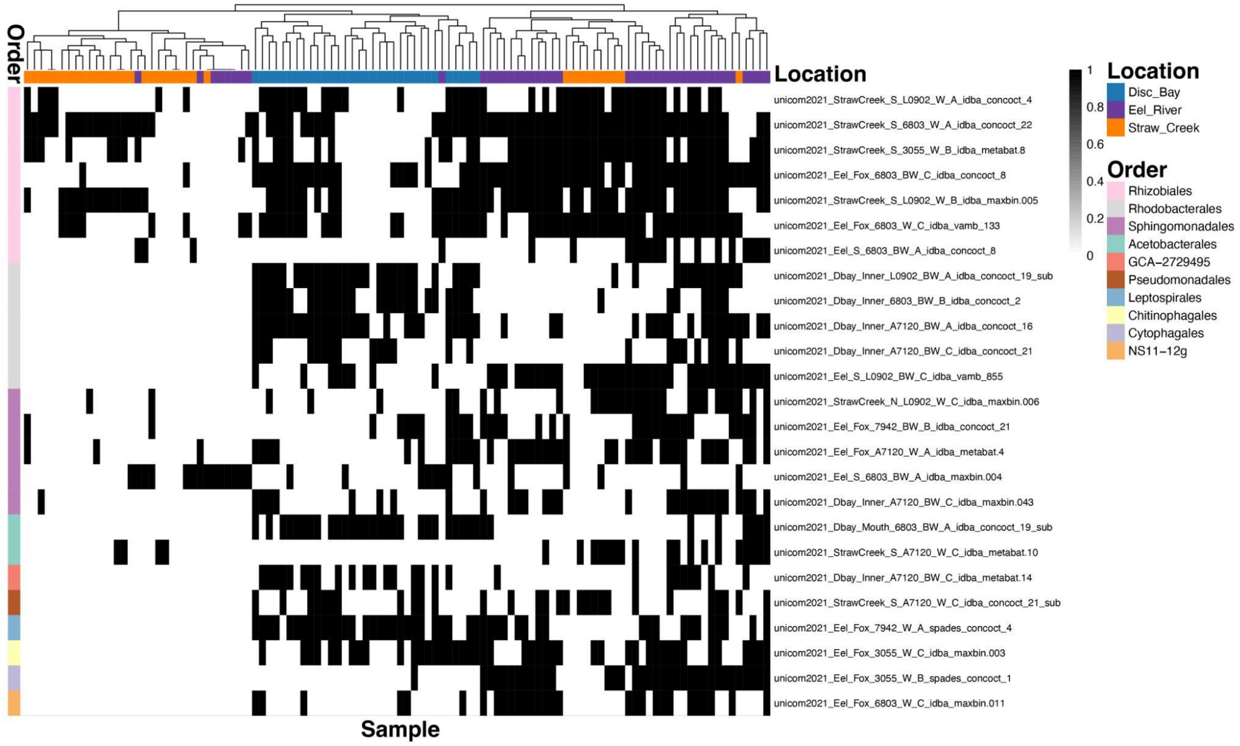


**Supplementary Fig. 6 | Heatmap of the distribution of core MAGs across samples.** The x-axis represents the samples, annotated with colored bars indicating the inoculum source location used for generating the respective co-cultures (Discovery Bay, Eel River, and Strawberry Creek). The y-axis represents core MAGs, annotated and ordered by their taxonomic classification at the order level, with the corresponding order indicated by the colored bar to the left of the heatmap. Presence or absence of a specific core MAG in each sample is depicted in binary (black for presence, white for absence). Samples (x-axis) are hierarchically clustered.

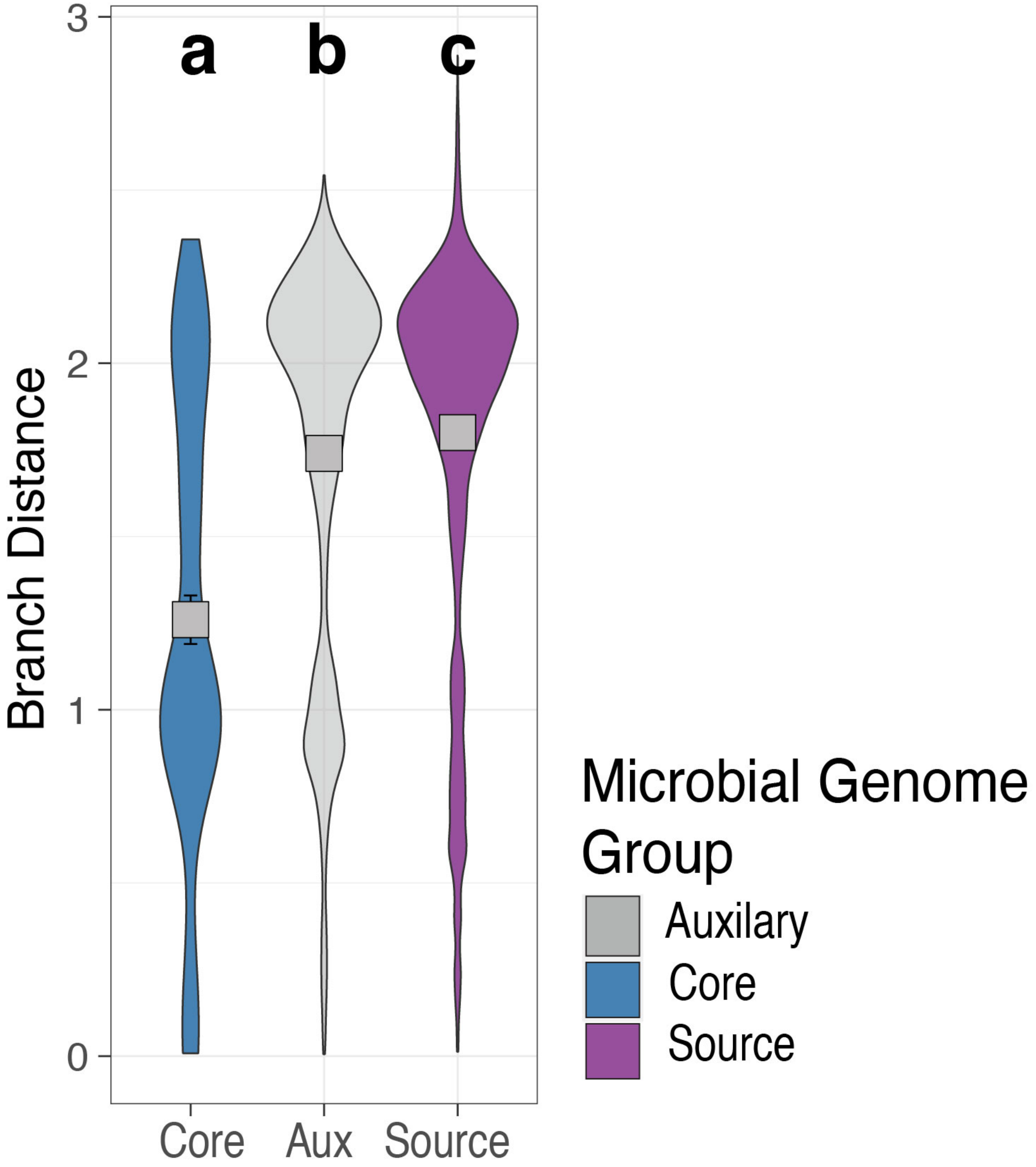


**Supplementary Fig. 7 | Distributions of pairwise all-vs-all phylogenetic branch distances.** Comparisons between MAGs within core (n = 300 comparisons), auxiliary (n = 43,017 comparisons), and source (n = 19,900 comparisons) groupings. Gray squares in the plot indicate the mean, and error bars indicate the 95% confidence interval. Statistical differences between groups are indicated by letters, and groups not sharing the same letter are significantly different (FDR ≤ 0.05; Pairwise Wilcoxon test).

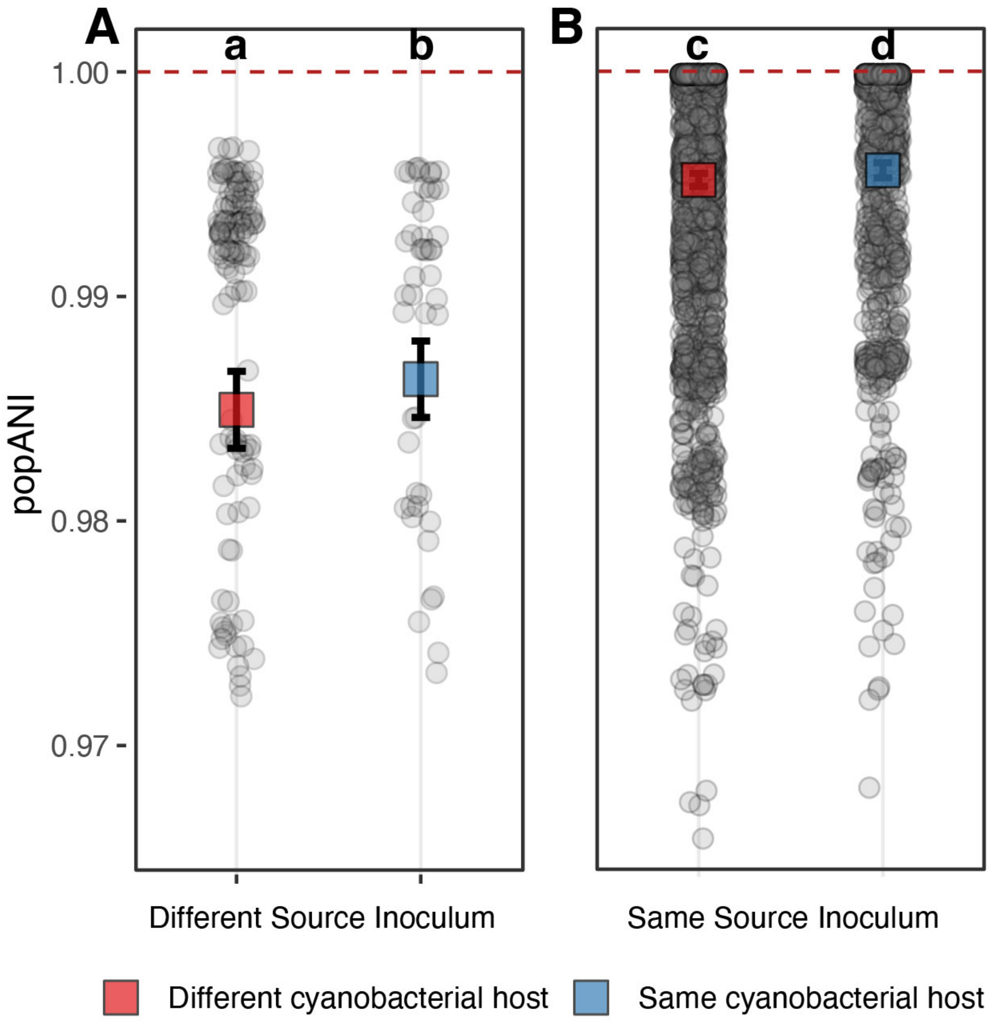


**Supplementary Fig. 8 | Population-level single nucleotide variant (SNV) analysis of microbial MAGs between samples.** The analysis involved 133 MAGs, including 13 core microbiome species, resulting in a total of 3088 pairwise comparisons (555 for core microbiome species). **(A)** Comparison of population average nucleotide identity (popANI) between *in vitro* community samples that have been inoculated using different source locations. For MAGs compared samples with different source inoculum no instances (0 out of 176 comparisons) of identical strain presence was detected. **(B)** Comparison of popANI between *in vitro* community samples that have been inoculated using the same source location. For MAGs compared between samples with the same source inoculum, 13.2% (386 out of 2912 comparisons) exhibited identical strains (popANI ≥ 99.999%). Beta regression analysis was utilized to understand the influence of shared cyanobacterial host strains and source inocula on popANI values. The results, indicated in both panels, demonstrate a significant relationship: shared source inocula significantly increased popANI values ($p < 0.0001$), while shared cyanobacterial host strains also had a notable impact ($p = 0.0073$). This suggests that both the source of inoculum and the cyanobacterial host strain play crucial roles in determining the genomic identity and similarity of microbial species within these communities. Square boxes in both plots represent estimated marginal means and whiskers indicate 95 % CI of estimated marginal means. Grey dots are the popANI values resulting from a single comparison of one MAG's SNPs between two samples. The red dotted line at the top of both plots indicates popANI = 99.999%. Letters above each column of a plot indicate statistically significant differences of estimated marginal means with columns not sharing a letter being statistically significantly different (FDR < 0.05).

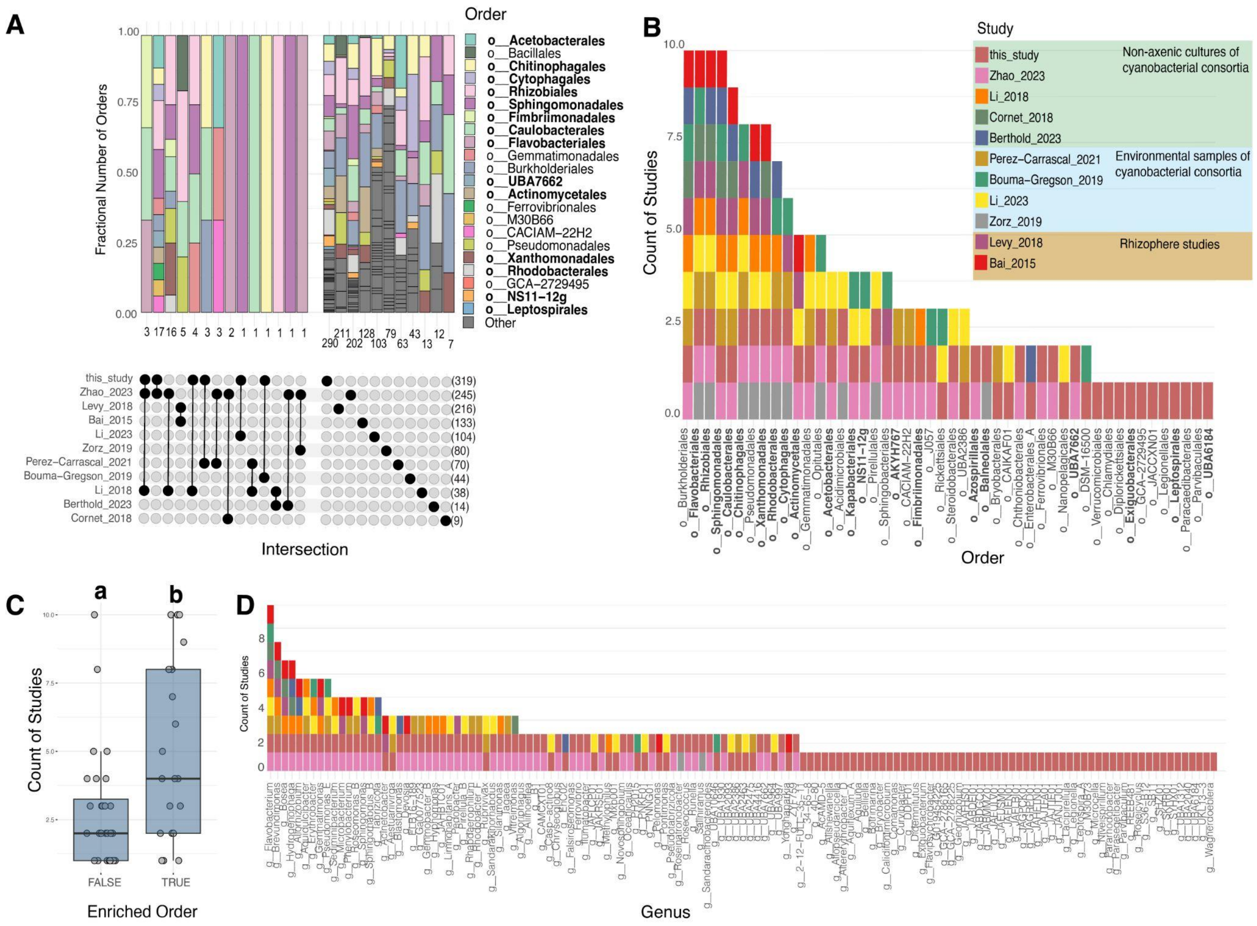


**Supplementary Fig. 9 | Shared taxa between different cyanobacterial and rhizosphere studies. (A)** Overlap of identical species between all 10 evaluated studies, and this study. Colored bars above UpSet plot in the left panel indicate the order-level taxonomy of overlapping species between studies. The number and order-level taxonomy of species unique to each study is shown in the right panel. Orders which were only observed in a single study were grouped into "Other" and colored gray. Numbers below the colored bars represent the total number of shared or unique species. Connected black dots in the UpSet plot below each bar illustrate the set of studies that share the species or that are unique for each study, with the numbers in parentheses indicating the total number of unique species for each examined study. **(B)** Number of studies in which each of the 48 order-level lineages identified in our study were also found. The bars represent the count of studies, with colors corresponding to different studies. Orders that are enriched in this study, as identified in **Fig. 2**, are shown in bold. **(C)** Comparison of the number of studies in which enriched orders were found. Box plots show the count of studies for orders that were not enriched (FALSE) versus those that were enriched (TRUE) during *in vitro* community establishment in this study. Letters above each boxplot indicate the groups are significantly different (p-value = 0.018; Wilcoxon Test). **(D)** Number of studies in which each of the 132 genus-level lineages identified in this study were also found. The bars represent the count of studies, with colors corresponding to different studies.

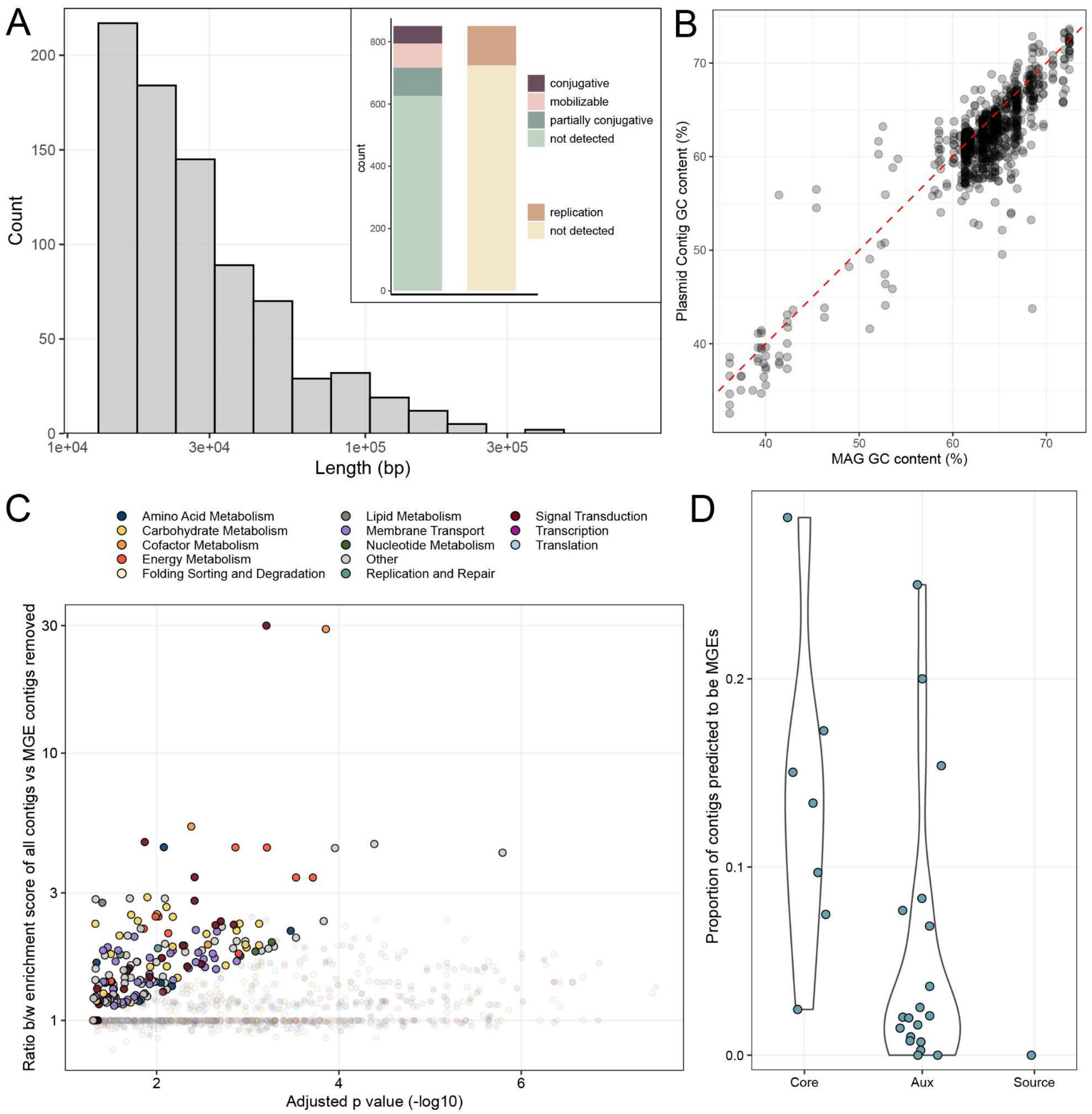


**Supplementary Fig. 10 | Features and functional analysis of putative plasmids recovered in this study. (A)** Distribution of all (n = 850) putative plasmid lengths across all analyzed MAGs. The inset bar plot shows the proportion of plasmids detected as conjugative, mobilizable, partially conjugative, and those with replication functions as well as the proportions where these functions were not detected. **(B)** Comparison of GC content between putative plasmids and their corresponding MAGs. The red dashed line represents a fitted regression line, indicating a significant positive correlation between the GC content of plasmids and their host MAGs. However, we note that while plasmid and MAG GC content are correlated, differential coverage binning enabled the association of plasmids with microbial MAGs even when GC content was divergent. **(C)** Enrichment analysis of KEGG Orthology (KO) groups on core microbiome plasmids versus non-core MAGs. Functional categories are color-coded, with significantly enriched KOs highlighted (FDR < 0.05). **(D)** Proportion of contigs predicted to be putative plasmids associated with *Rhizobiales* MAGs of the core microbiome, auxiliary MAGs, and MAGs from source environment samples.

Core microbiome *Rhizobiales* MAGs had a significantly higher proportion of contigs predicted to be putative plasmids compared to auxiliary species and source environment MAGs.

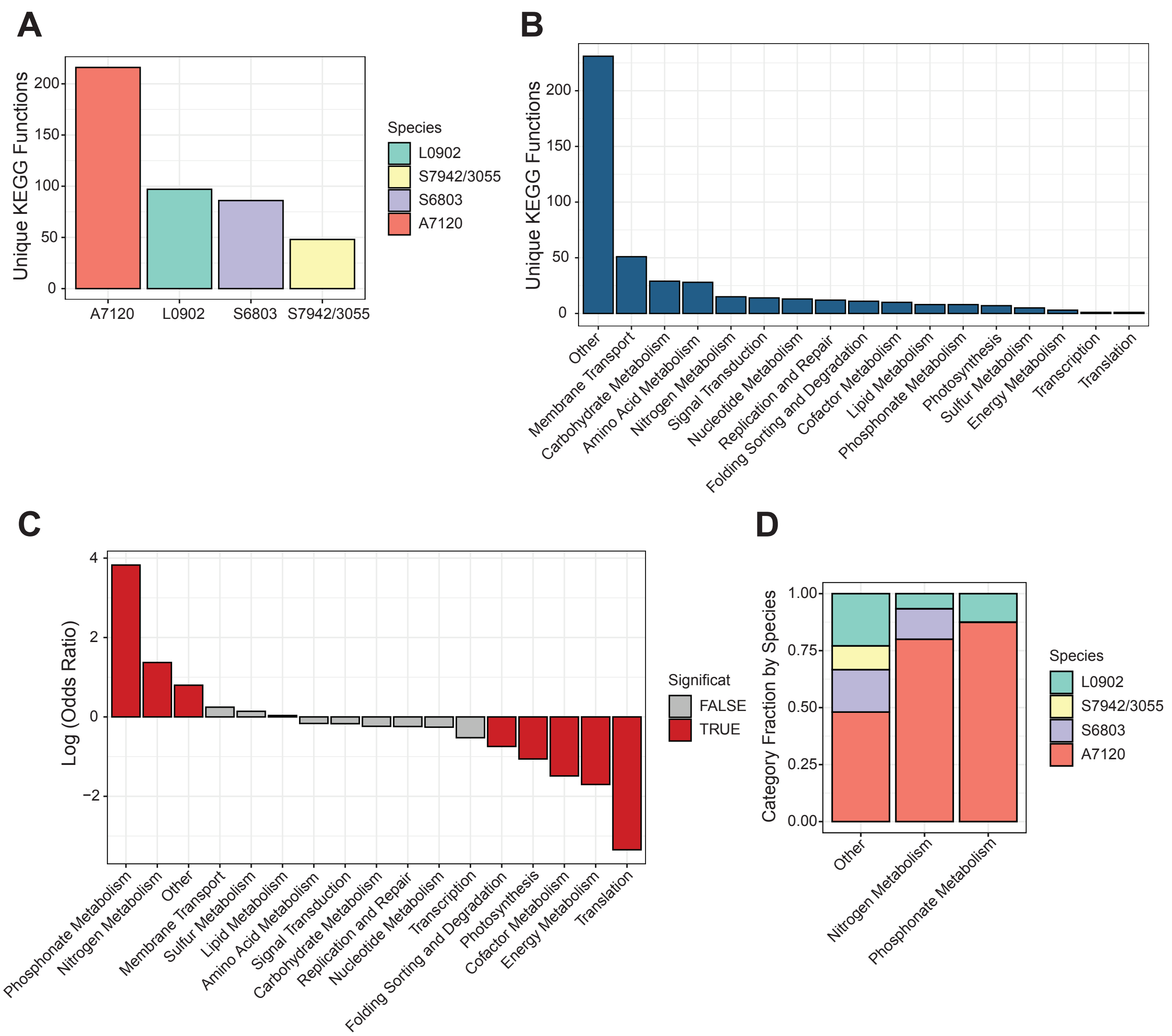


**Supplementary Fig. 11 | Unique KEGG functions across the cyanobacterial host species used in this study. (A)** Total number of KEGG KOs that were unique to each host species genome evaluated in the study. *Synecococcus elongatus PCC 7942* and *3055* are strain variants and were only represented by one species-representative genome in our analysis. **(B)** Total number of KEGG KOs that were unique across all host species broken down by high-level functional categories. **(C)** Differential enrichment of unique KOs (e.g. identified in only one host species genome) relative to non-unique KOs (e.g. identified in more than one host species genome) for each functional category across all host species. The X-axis indicates the log transformed Fisher odds ratio with positive values indicating the functional category had significantly more unique KOs associated with it than by chance, and negative values indicating the functional category had significantly more unique KOs associated with it than by chance. Functional categories where over or under-enrichment of unique KOs was significant (FDR < 0.05; Fisher Exact Test) have red colored bars. **(D)** Fractions of unique KOs broken down by the fraction found in each host species for the three functional categories found to be over-enriched in unique KOs across all host species. Also see **Tables S24** and **S25**.
